# Generation of floxed alleles for cell-specific knockout in zebrafish

**DOI:** 10.1101/2022.09.06.506849

**Authors:** Masahiro Shin, Takayuki Nozaki, Benjamin Toles, Amy Kolb, Kevin Luk, Sumio Isogai, Kinji Ishida, Tomohito Hanasaka, Michael J. Parsons, Scot A. Wolfe, Nathan D. Lawson

**Affiliations:** Department of Molecular, Cell, and Cancer Biology, University of Massachusetts Chan Medical School, Worcester, MA 01605, USA; Technical Support Center for Life Science Research, Iwate Medical University, Shiwa, Iwate 028-3694, Japan; Department of Medical Education, Iwate Medical University, Shiwa, Iwate 028-3694, Japan; Department of Developmental and Cell Biology, School of Biological Sciences, University of California, Irvine, Irvine, CA 92697, USA

## Abstract

A benefit of the zebrafish as a model is its amenability to genetic approaches. However, a lack of conditional deletion alleles makes it a challenge to perform stage- or cell-specific knockout. Here, we initially applied an existing protocol to establish a floxed allele for *gata2a* and encountered several issues, including off-target integration and incomplete knock-in. To address these problems, we developed a protocol incorporating simultaneous co-targeting with Cas12a to insert loxP sites in *cis*, together with transgenic counter-screening to identify off-target insertions. We applied this protocol to establish a floxed allele of *foxc1a* in a single generation. We demonstrate the utility of our floxed alleles by verifying Cre-dependent deletion, which yielded expected phenotypes in each case. Finally, we used the floxed *gata2a* allele to demonstrate an endothelial autonomous requirement in lymphatic valve development. Together, our results provide a framework for straightforward generation and application of floxed alleles in zebrafish.

## INTRODUCTION

Over the past several decades, the zebrafish has become a widely accepted model system for studying developmental biology. Several notable benefits of the zebrafish life cycle have contributed to its success in this regard. Zebrafish embryos are transparent and develop externally, allowing detailed visualization of morphogenesis *in vivo* (Beis and Stainier, 2006). Embryogenesis is highly synchronous and rapid, with most organ systems functioning by one to two days post fertilization (dpf) (Kimmel et al., 1995). Coupled with these benefits is the amenability of the zebrafish to genetic approaches. Zebrafish adults are easy to maintain and exhibit high fecundity, with a single pair of adults often yielding hundreds of eggs on a given day. Zebrafish researchers initially leveraged these characteristics to perform large-scale forward genetic screens for a host of embryonic phenotypes (Driever et al., 1996; Haffter et al., 1996; Mullins et al., 2021). Subsequent development of sequence-specific nucleases, such as those that target clustered regularly interspaced short palindromic repeats (CRISPR) in bacterial genomes, has allowed introduction of targeted germline deletions in the zebrafish genome for reverse genetic approaches (Hruscha et al., 2013; Hwang et al., 2013). Thus, a combination of genetic accessibility and ideal embryonic characteristics make the zebrafish as ideal model for interrogating gene function during development.

Nearly all zebrafish mutants perturb the function of a given gene from the earliest point at which it is expressed during development, making it a challenge to study direct roles in subsequent processes. While mosaic analysis through cell transplantation can address issues of cell autonomy in these mutants, ascribing primary functional effects at later stages remains challenging. Early efforts to address this issue relied on identification of temperature sensitive alleles, but these were limited (Johnson and Weston, 1995). More recently, conditional transgenic systems have been developed, but these rely on dominant activating or inhibitory transgenes that may interfere with unrelated pathways (Carney and Mosimann, 2018). Thus, a reliable and definitive conditional genetics platform is lacking in zebrafish, limiting genetic analysis at postembryonic and adult stages.

The mouse became an established genetic model through development of techniques for making germline deletions using embryonic stem cells (Doetschman et al., 1987; Mansour et al., 1988; Thomas and Capecchi, 1987). Subsequently, mouse researchers leveraged this platform for conditional genetics using the Cre/lox system (Gu et al., 1994). Cre is a bacteriophage recombinase that stimulates recombination between loxP sequences (Nagy, 2000). When loxP sites are arranged directly in *cis*, recombination results in deletion of the intervening sequence. By modifying endogenous mouse loci to flank coding exons with loxP sites (referred to as “floxed”), together with transgenic expression of a cell-specific Cre, it is possible to achieve tissue-specific gene knockout (Gu *et al*., 1994). Timing of the knockout can be controlled using Cre fused to a modified estrogen ligand binding domain that binds to tamoxifen (CreERT; (Feil et al., 1996)). To date, there are over 3000 published floxed alleles for approximately 2300 mouse genes, along with more than 300 transgenic lines expressing CreERT, allowing comprehensive genetic analysis in any biological context.

In zebrafish, the Cre/lox system is functional and has been applied for lineage tracing and conditional transgene expression (Carney and Mosimann, 2018). However, using Cre/lox for conditional gene deletion in zebrafish has been limited by the difficulty in generating floxed alleles. Indeed, only two previous studies have successfully such alleles in zebrafish. Burg et al. used single-stranded oligodeoxynucleotides (ssODNs) with short homology arm sequences and a loxP site as a template for homology-directed repair (HDR), along with targeting an endogenous site using Cas9 nuclease, to generate a floxed *tbx20* allele (Burg et al., 2018). However, this approach required each loxP site to be inserted sequentially, requiring at least two generations to establish a stable floxed allele. By contrast, Hoshijima et al. relied on a plasmid template with 1kb homology arms to replace an endogenous exon of *potassium voltage-gated channel, subfamily H, member 6a* (*kcnh6a*) with a floxed version in a single generation (Hoshijima et al., 2016). In both cases, Cre-mediated excision of the floxed allele resulted in expected cardiac defects (Burg *et al*., 2018; Hoshijima *et al*., 2016). Here, we present our experience in applying the Hoshijima et al. technique to generate a floxed *gata2a* allele, with a practical emphasis on pitfalls and points for improvement. We subsequently leveraged our findings to develop an improved knock-in protocol, which we demonstrate through generation of a floxed allele for *foxc1a*, a single exon gene. Finally, we used the *gata2a* floxed allele together with a tissue-specific inducible CreERT transgene to demonstrate an endothelial autonomous requirement during lymphatic valve development.

## RESULTS

### Generation of a *gata2a* conditional knock-in allele

To generate a floxed allele we chose the Hoshijima protocol (Hoshijima *et al*., 2016) since it should yield knock-in founders in a single generation. As a target, we chose *gata2a*, which encodes a zinc finger transcription factor expressed in multiple cell types, including blood, endothelial cells, and neurons (Andrzejczuk et al., 2018; Pase et al., 2009; Shin et al., 2019). Loss of *gata2a* results in circulatory defects, improper spinal neuron specification, lymphatic valve abnormalities, and failure to inflate the swim bladder (Andrzejczuk *et al*., 2018; Shin *et al*., 2019; Zhu et al., 2011). To generate a floxed *gata2a* allele, we constructed a plasmid with homology arms spanning 1 kb up- and downstream of exon 5, which was flanked by loxP sites (**Figure 1A, B**). Exon 5 deletion results in an exon 4/6 junction that remains in-frame, but removes the C-terminal zinc finger (**Supplementary Figure S1A, B**), which is essential for DNA binding in GATA proteins (Martin and Orkin, 1990). Our targeting strategy is similar to that applied to generate the widely used mouse Gata2^tm1Sac^ allele (Charles et al., 2006), which results in the same in-frame sequence following deletion (**Supplementary Figure S1C**).

**Figure 1.**
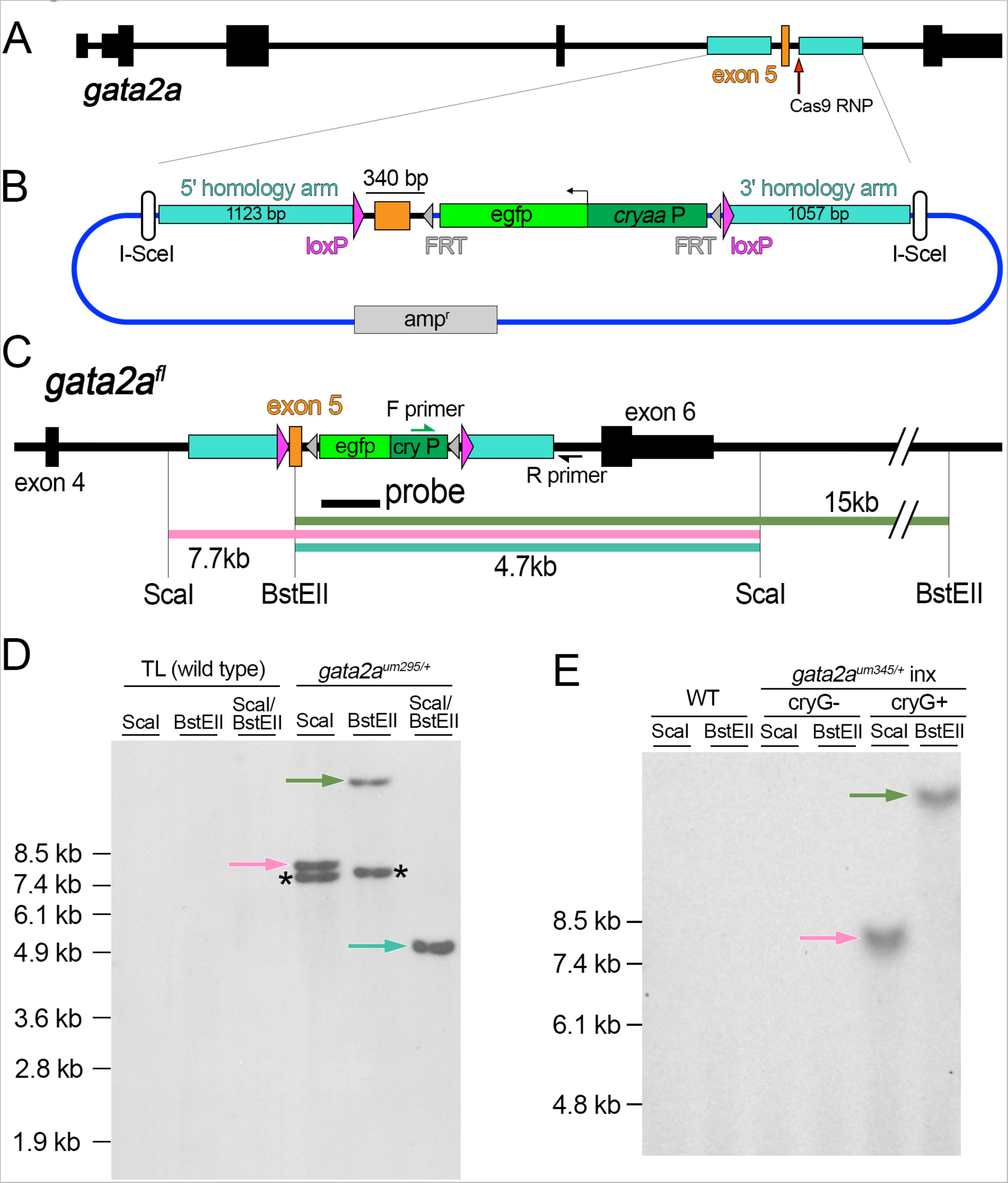
Generation of a *gata2a* conditional deletion allele. (A) Zebrafish *gata2a* locus highlighting exon 5, homology arms, and Cas9 ribonucleoprotein (RNP) target. (B) Targeting plasmid used to generate a floxed *gata2a* allele. egfp – enhanced green fluorescent protein, cryaa:P – cryaa promoter element, *amp^r^* – ampicillin resistance cassette. (C) Floxed *gata2a* allele (*gata2a^fl^*). Expected fragment sizes following restriction digest of genomic DNA with indicated enzyme are shown. (D, E) Southern analysis using genomic DNA from embryos of indicated genotype. Blots were hybridized with a DIG-labeled *egfp* probe and detected by chemiluminescence. Arrows denote fragments of expected size in the event of precise integration. Asterisk denotes fragments resulting from off-target integration.

In the targeting construct, we used homology arm sequences that matched the haplotype in adult fish used to generate embryos for injection, as recommended previously (Hoshijima *et al*., 2016). Downstream of exon 5, we placed a cassette comprising enhanced green fluorescent (EGFP) driven by the *crystallin, alpha A* (*cryaa*) promoter in an orientation opposite to the endogenous locus (**Figure 1B**; **Supplementary File S1**). The *cryaa* promoter drives Egfp expression in the lens and allows identification of individuals carrying the transgene (Kurita et al., 2003), although it does not distinguish on- and off-target insertions. The *cryaa:egfp* cassette is flanked by FRT sites to allow for removal using FLP recombinase in the event that it interferes with normal gene function. The entire targeting cassette was flanked by sites for the I-SceI nuclease for release of the linear HDR template (Hoshijima *et al*., 2016).

To stimulate HDR, we used a Cas9/sgRNA ribonucleoprotein complex (RNP) to introduce a double strand break (DSB) at a sequence immediately downstream of *gata2a* exon 5 (**Figure 1A**). In the targeting construct, we disrupted the Cas9 spacer sequence with the 3’ loxP site to prevent cleavage of the targeting construct or the inserted transgene (**Supplementary File 1**). We co-injected 1-cell stage embryos with Cas9 RNP, circular plasmid targeting construct, and I-SceI. At 3 days post fertilization (dpf), PCR across the 3’ homology arm junction revealed 6 out of 12 individual embryos that exhibited a fragment of correct size and junction sequence in *cryaa:egfp*-positive embryos (**Figure S2A-C**). We repeated injections, grew *cryaa:egfp*-positive embryos to adulthood, and identified 12 founders out of 115 that transmitted *cryaa:egfp* to progeny embryos. Of these, only one gave PCR-positive embryos with the primers above, suggesting that most founders carried off-target insertions. The allele with evidence for targeted knock-in is hereafter referred to as *gata2a^um295^*.

Subsequent analysis of *gata2a^um295^* revealed two issues. First, only the 3’ loxP site and *cryaa:egfp* cassette were inserted at the target (**Supplementary Figure S2D-F**). Second, Southern analysis revealed an off-target integration. If precise knock-in occurs, we expect single fragments of 7.7kb or 15kb with digests of *gata2a^um295^* genomic DNA using the restriction enzymes ScaI or BstEII, respectively, while cutting with both yields a 4.7kb fragment (**Figure 1C**). Southern blotting of digested *gata2a^um295^* genomic DNA showed fragments of these approximate sizes when hybridized with an Egfp probe, consistent with integration at the targeted site (**Figure 1D**). However, we noted a second band in ScaI- and BstEII-digested samples, and a possible 4.7 kb doublet from double digests (**Figure 1D, extra bands marked by asterisks**), which could arise from integration of vector backbone cut only once by I-SceI immediately following injection (for example, see **Supplementary Figure S3A**). Unfortunately, this off-target integration also co-segregated with *gata2a^um295^* indicating that it was linked to *gata2a*. Since the linked off-target insertion likely retained both loxP sites (**Supplementary Figure S3A**), it could lead to large chromosome deletions or inversions in the presence of Cre. To test this possibility, we injected *cre* mRNA into 1-cell stage embryos from *gata2a^um295^* carriers and performed Southern analysis. As above, two Egfp-positive fragments from genomic DNA digested with ScaI or BstEII were evident in uninjected *gata2a^um295^* embryos (**Supplementary Figure S3A-C**). By contrast, following injection with *cre* mRNA we only identified single bands consistent with the correctly targeted insert, which, as noted, lacks a 5’ loxP site. Thus, recombination between the off-target loxP sites resulted in deletion of the off-target exon5/*cryaa:egfp* cassette (**Supplementary Figure S3A-C**). Importantly, we did not observe any additional shifts in fragment sizes suggesting that no recombination had occurred between the on- and off-target loxP sites.

Since we did not detect Cre-stimulated recombination between on- and off-target sites in *gata2a^um295^*embryos, we proceeded to introduce a 5’ loxP site upstream of the targeted exon 5. We first injected *gata2a^um295^* embryos with *cre* mRNA to eliminate the off-target *cryaa:egfp* cassette and grew these to adulthood (referred to as *gata2a^um329^*). We would note that *gata2a^um329^* fish still bear residual vector backbone and a single loxP sequence at the off-target site. We subsequently injected *gata2a^um329^* embryos with a Cas12a RNP targeted to sequence upstream of exon 5, along with a double-stranded (ds) ODN template with 30 bp flanking homology arms and a loxP site in direct orientation relative to the integrated 3’ loxP site (**Supplementary Figure S3D, E**). We identified a founder with a precise 5’ loxP integration (**Supplementary Figure S3F**) and the resulting floxed allele (*gata2a^um345^*) is hereafter referred to as *gata2a^fl^* (**Figure 1C,E**).

To test *gata2a^fl^* functionality, we crossed *gata2a^+/fl^*carriers to *gata2a^+/um27^* adults, which bear a frameshift deletion upstream of the Gata2a zinc finger domains (Zhu *et al*., 2011). As expected, one-half of progeny embryos showed *cryaa:egfp* expression and none of these show evidence of exon 5 deletion in the absence of Cre (**Figure 2A, B**; **Supplementary Table S1**). By contrast, *cre* mRNA injection caused loss of *cryaa:egfp* expression in all embryos and deletion of exon 5 in approximately one-half of embryos (**Figure 2B, Supplementary Table S1**). We also observed fully penetrant loss of swim bladder inflation in heterozygous *gata2a^um27^* embryos that exhibited Cre-mediated exon 5 deletion (*gata2a^um27/Δ^*), while sibling embryos were normal (**Figure 2C, Supplementary Table S1**). Uninjected trans-heterozygous *gata2a^um27/fl^*larvae inflated their swim bladders at 5 dpf and otherwise appeared indistinguishable from *gata2a^+/fl^* siblings (**Figure 2D**, see below). Consistently, *gata2a^fl/fl^*adults were viable and fertile. Similar to the *gata2a^um27^* complementation cross, uninjected *gata2a^fl/fl^* larvae from crosses between *gata2a^fl/fl^* and *gata2a^+/fl^* adults expressed *cryaa:egfp* and displayed normal swim bladder inflation at 6 dpf (**Figure 2E, Supplementary Table S2**). By contrast, *gata2a^fl/fl^* larvae injected with a zebrafish codon-optimized *cre* mRNA (*zfcre*) at the 1-cell stage showed loss of *cryaa:egfp* and failed to inflate their swim bladders by 6 dpf (**Figure 2F, Supplementary Table S2**).

**Figure 2.**
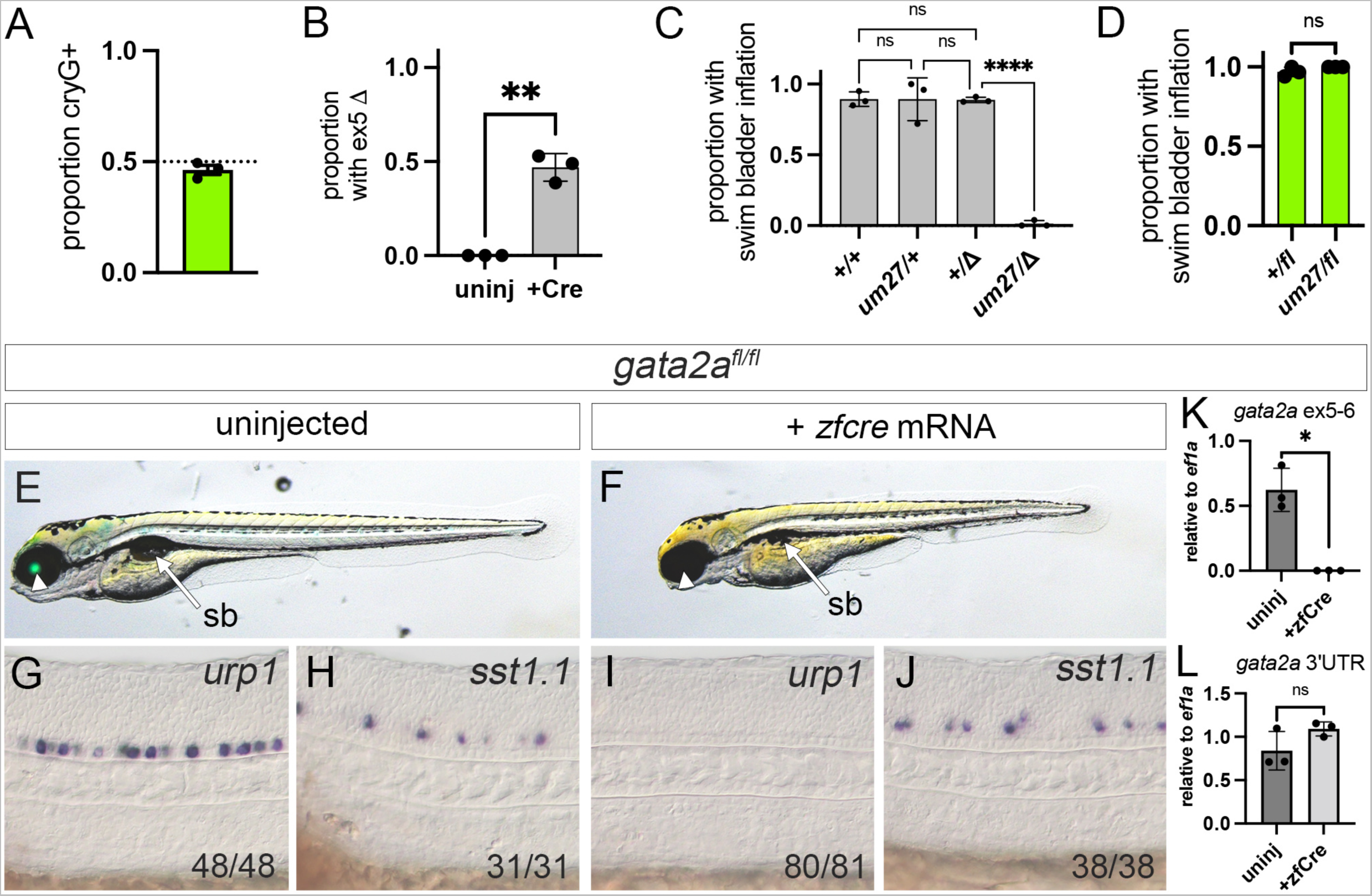
Functional validation of a floxed *gata2a* allele. (A-D) Embryos from crosses between *gata2a^+/um27^*and *gata2a^+/fl^* individuals (n = 3 clutches). (A) Proportion of embryos with lens *Egfp* (cryG+) expression at 2 dpf. (B) Proportion of uninjected or *cre* mRNA-injected embryos with exon 5 deletion. ** p<0.01, paired t-test. (C) Proportion of embryos of indicated genotype with inflated swim bladder following injection with *cre* mRNA. “Δ” denotes deletion of exon 5 confirmed by PCR. Analysis of variance, p<0.0001. **** p<0.0001, Tukey’s multiple comparison test. (D) Proportion of uninjected *cryaa:egfp*-positive embryos of indicated genotype with inflated swim bladder at 5 dpf. ns – not significant, Wilcoxon test. (E-J) Homozygous *gata2a^fl/fl^* larvae or embryos. Lateral views, anterior to the left, dorsal is up. (E, G, H) Uninjected embryos. (F, I, J) Embryos injected with *zf-cre* mRNA at 1-cell stage. (E, F) Overlay of transmitted light and green fluorescence at 5 dpf. Arrowhead denotes lens and Egfp expression, or absence thereof, arrow indicates swim bladder (sb). (G-J) Differential interference contrast (DIC) images of 24 hpf embryos subjected to whole mount in situ hybridization with antisense riboprobes against indicated transcript. (K, L) Quantitative RT-PCR of uninjected or *zf-cre* mRNA-injected *gata2a^fl/fl^* embryos/larvae at 24 hpf using primers to detect *gata2a* (K) exon 5 and 6, or (L) 3’ UTR. Paired t-test, * - p<0.05, ns – not statistically significant.

We next assessed spinal neuron specification and trunk circulatory function, both of which are affected in *gata2a^um27^* embryos (Andrzejczuk *et al*., 2018; Zhu *et al*., 2011). In uninjected *gata2a^fl/fl^*embryos, we detected *urotensin related peptide 1* (*urp1*) transcript in the ventral-most Kolmer-Agduhr (KA) neurons in the floor plate and *somatostatin 1, tandem duplicate 1* (*sst1.1*) in more dorsally-located KA neurons at 24 hpf (**Figure 2G, H;** (Andrzejczuk *et al*., 2018)). By contrast, *urp1*-positive cells were absent in *gata2a^fl/fl^* embryos injected with *zf-cre* mRNA, while *sst1.1*-expressing KA neurons were not affected (**Figure 2I, J**). This defect is identical to that seen in *gata2a^um27^* mutant embryos (Andrzejczuk *et al*., 2018). We also observed circulatory defects in *gata2a^fl/fl^*. Control *gata2a^+/fl^* sibling embryos injected with *zf-cre* mRNA exhibited a heartbeat and circulation throughout aortic arch blood vessels, along with venous circulatory return from cranial vessels through the posterior cerebral vein and primary head sinus (**Supplementary Movie S1**). Trunk circulation through the dorsal aorta (DA) and posterior cardinal vein (PCV) was also normal (**Supplementary Movie S1**). In zfCre-injected *gata2a^fl/fl^*embryos, we similarly observed a heartbeat along with blood cell circulation through aortic arch blood vessels and cranial veins, albeit weaker than in *gata2a^+/fl^* siblings (**Supplementary Movie S2**). However, DA and PCV circulation was absent (**Supplementary Movie S2**), similar to *gata2a^um27^* mutants (Zhu *et al*., 2011).

Analysis of *gata2a* transcript in *gata2a^fl/fl^* embryos by qRT-PCR using primers in exon 5 and 6 indicated expression in uninjected embryos, but not in siblings injected with *zfcre* mRNA (**Figure 2K**). By contrast, *gata2a* transcript detected using 3’ UTR primers was not reduced by deletion of exon 5, consistent with an in-frame exon 4/6 fusion (**Figure 2L, Supplementary Figure S1B**). Taken together, these observations indicate that *gata2a^fl^* provides wild type *gata2a* function and Cre-dependent deletion of exon 5 in *gata2a^fl/fl^* phenocopies *gata2a^um27^*mutants. Thus, *gata2a^fl/fl^* behaves as a conditional loss-of-function allele.

### Generation of a *foxc1a* conditional allele using an improved knock-in protocol

Following our experience with the *gata2a^fl^* allele, we sought to make improvements that would promote complete knock-in of a loxP-flanked cassette and would allow easier identification of off-target insertions. As a target, we chose *foxc1a*, a single exon gene that encodes a widely expressed Forkhead transcription factor (Topczewska et al., 2001). *Foxc1a* mutant embryos display numerous phenotypes, including those associated with cardiovascular, somite, and jaw development (Banerjee et al., 2015:Ferre-Fernandez, 2020 #937; Li et al., 2015; Shin *et al*., 2019; Whitesell et al., 2019). For most phenotypes, a cell autonomous requirement for *foxc1a* has not been demonstrated in zebrafish. Thus, a conditional floxed allele would be valuable to better interrogate the cellular nature of *foxc1a* deficiency.

A problematic issue at the *gata2a* locus was incomplete knock-in leading to a failure to insert the upstream loxP site. We hypothesized that 340 bp upstream of the Cas9 target in the *gata2a* gene was sufficient to allow HDR-mediated integration of the downstream cassette without inserting the 5’ loxP site (**see Figure 1A**). To avoid this issue, we applied simultaneous CRISPR targeting at both insertion points for loxP sites flanking the *foxc1a* coding sequence (**Figure 3A, B**). In this case, we used Cas12a as a nuclease since it can stimulate higher HDR rates than Cas9 in mammalian cells and zebrafish embryos (Moreno-Mateos et al., 2017; Shahbazi et al., 2019). Within the *foxc1a* targeting construct, we included homology arms extending 400 bp upstream and 1kb downstream of the respective Cas12a sites and placed loxP sites at or near these targets (**Figure 3B**; **Supplementary File 2**). As we did for *gata2a*, we matched homology arm sequences to the haplotype of embryos used for injection. To detect off-target integrations, we used a vector where *mcherry* was driven by the cardiomyocyte-specific *myosin, light chain 7* (*myl7*) promoter (Hoshijima *et al*., 2016; Huang et al., 2003). The *myl7:mcherry* cassette lies outside the homology arms, but within the I-SceI sites (**Figure 3B, Supplementary File S2**), allowing identification of individuals carrying off-target insertions by virtue of mCherry expression in the heart.

**Figure 3.**
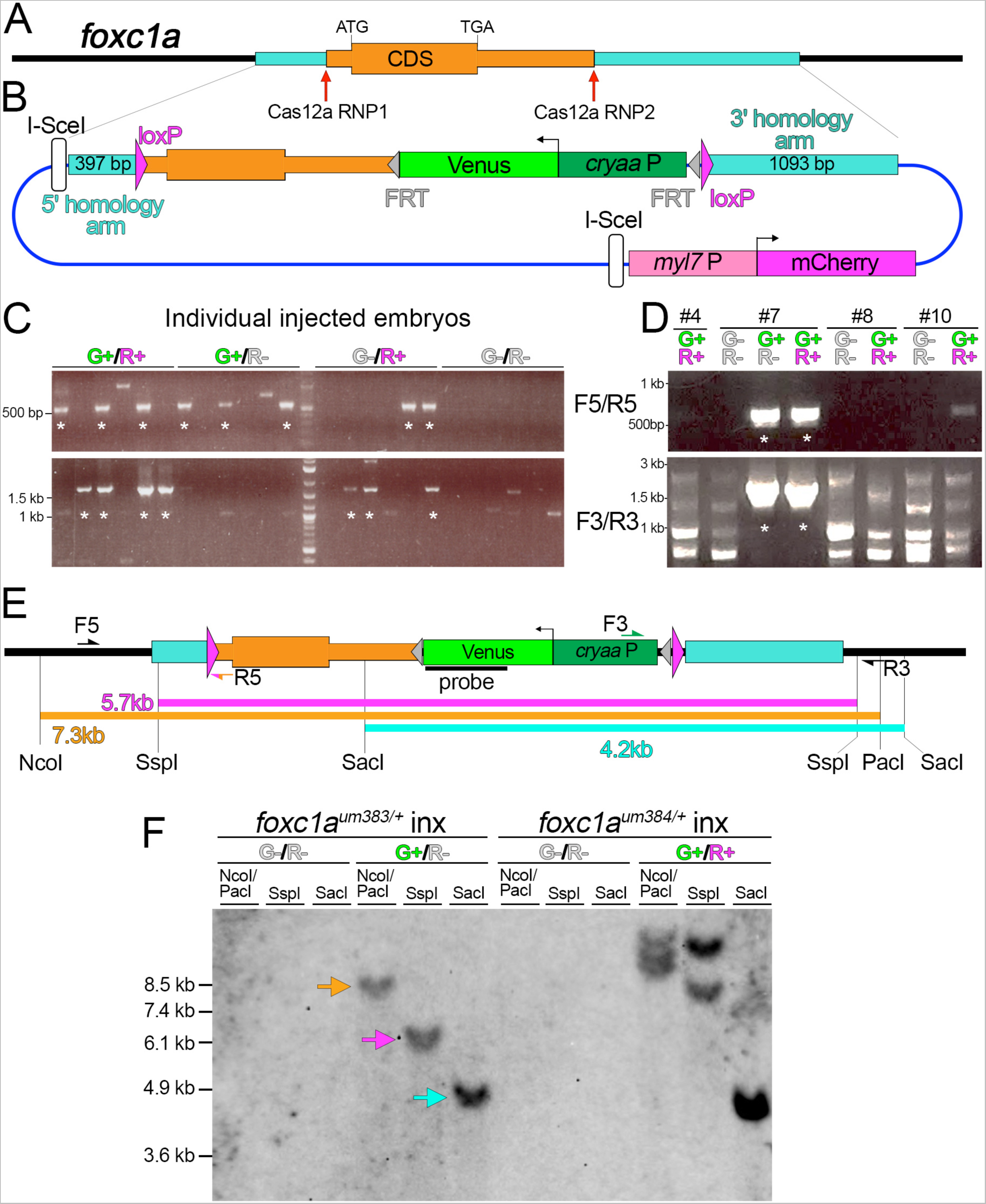
Generation of a *foxc1a* conditional deletion allele. (A) Zebrafish *foxc1a* locus with location homology arms highlighted (turquoise) and Cas12a ribonucleoprotein (RNP) targets indicated. Coding sequence (CDS), bounded by start (ATG) and stop codons (TGA) is indicated. (B) Targeting plasmid used to generate a floxed *foxc1a* allele. *cryaaP* – *cryaa* promoter. Schematic is not to scale as the *myl7:mcherry* cassette is immediately adjacent to the 3’ homology arm; see Supplementary File S2. (C, D) PCR of 3 dpf embryos scored for *cryaa:venus* (G+) and *myl7:mcherry* (R+) expression with indicated primer pairs (F5/R5 or F3/R3), location of which are shown in (E). Asterisk denotes lanes with product of expected size. (C) Individual embryos following injection with Cas12a RNPs, targeting construct, and I-SceI. (D) Pooled F1 embryos from 4 putative P0 founders. (E) Floxed *foxc1a* allele (*foxc1a^fl^*). Expected fragment sizes following restriction digest with indicated enzymes are shown. (F) Southern analysis using genomic DNA from embryos of indicated genotype and screened for expression of Venus (“G”) or mCherry (“R”). Blots were hybridized with a DIG-labeled *venus* probe and detected by chemiluminescence.

We co-injected both Cas12a RNPs, along with targeting construct, and I-SceI into 1-cell stage wild type embryos bearing the matching *foxc1a* haplotype sequence. At 2 dpf, we separated embryos by *cryaa:venus* and *myl7:mcherry* expression or co-expression and assessed knock-in by PCR across both homology arm junctions using anchored primers specific to the target construct (**Figure 3C, E**). Approximately one-third of the injected embryos exhibited co-expression of *cryaa:venus* and *myl7:mcherry* (19 out of 52) and the same proportion with only *cryaa:venus* expression (19 out of 52). A smaller proportion exhibited only *myl7:mcherry* or no transgene expression (7 out of 52 for each class). Surprisingly, only embryos co-expressing both *cryaa:venus* and *myl7:mcherry* consistently showed evidence of complete targeted knock-in, while those expressing only *cryaa:venus* did not (**Figure 3C**). Thus, the *myl7:mcherry* cassette does not distinguish on-versus off-target integration at this stage.

We repeated co-injections and grew embryos that co-expressed *cryaa:venus* and *myl7:mcherry* to adulthood, at which point they were subjected to individual incrosses with wild type adults to identify founders. From 45 putative founders (P0), we identified 9 fish that gave progeny embryos with *cryaa:venus* expression (**Supplementary Table S3**). Seven of nine P0 fish gave rise only to embryos that were double-positive for *cryaa:venus* and *myl7:mcherry*, with all but one being negative following PCR across 5’ and 3’ junctions (for examples see embryo pools #4, 8, and 10 in **Figure 3D, Supplementary Table S3**), indicating that they bore off-target insertions. In the single double-positive founder with evidence of knock-in, *myl7:mcherry* was not genetically separable from *cryaa:venus* suggesting that NHEJ-mediated insertion of an additional transgene copy occurred at the *foxc1a* locus. We also identified 2 founders that transmitted a mix of *cryaa:venus*;*myl7:mcherry* double-positive and *cryaa:venus* single-positive sibling embryos (**Supplementary Table 3**), one of which gave embryos with evidence of complete knock-in by PCR (**Figure 3D, founder #7**). Interestingly, both double-positive and single-positive embryos in this case were positive for 5’ and 3’ junction PCRs (**Figure 3D**).

The results above suggested that founder #7 carried at least two distinct insertions, one of which was the desired knock-in at the *foxc1a* locus. To confirm this, we separately grew *cryaa:venus*-positive;*myl7:mcherry-negative* and *cryaa:venus*;*myl7:mcherry* double-positive sibling embryos to adulthood. These alleles are referred to as *foxc1a^um383^* and *foxc1a^um384^*, respectively. At adulthood, we in-crossed individual *foxc1a^+/um383^* or *foxc1a^+/um384^* carriers and separated the progeny based on transgene expression, followed by Southern blot analysis. Precise HDR-mediated knock-in at the target site would yield single fragments of approximately 7.3kb with a double digest of genomic DNA using NcoI and PacI, and 5.7kb or 4.2kb following single digests with SspI or SacI, respectively (**Figure 3E**). In *cryaa:venus*-positive; *myl7:mcherry-negative* embryos from *foxc1a^um383^*parents, we observed fragments of approximate sizes consistent with precise knock-in at the target site, while genomic DNA from *cryaa:venus-negative* embryos did not hybridize to the Venus probe (**Figure 3E, F**). By contrast, we observed multiple fragments, none of which were of the expected size, in digested genomic DNA of double-positive *cryaa:venus;myl7:mcherry* embryos from *foxc1a^um384^* carriers (**Figure 3F**). These observations confirm that multiple germline insertions independently occurred in founder #7: *foxc1a^um383^* is a precisely targeted single copy knock-in allele, referred to hereafter as *foxc1a^fl^*, while *foxc1a^um384^* bears multiple off-target insertions. Importantly, our results demonstrate that counter-screening using the *cryaa:venus* and *myl7:mcherry* markers can allow preliminary identification of F1 progeny that bear putative on- or off-target insertions, respectively.

To confirm the functionality of the *foxc1a^fl^* allele, we crossed carriers to heterozygous *foxc1a^p162^* adults, which bear a nonsense mutation in the Forkhead domain (Banerjee *et al*., 2015). All embryos resulting from this cross, including *foxc1a^p162/fl^* transheterozygotes, appeared with normal morphology (**Figure 4A**), indicating that the *cryaa:venus* cassette does not affect *foxc1a* function (**Figure 4A, bottom panel**). In sibling embryos injected with *cre* mRNA, we observed a complete loss of *cryaa:venus* expression (**Figure 4B**). Furthermore, transheterozygous *foxc1a^p162/fl^* embryos injected with *cre* mRNA showed deletion of the *foxc1a* conditional allele (*foxc1a^p162/Δ^*) and exhibited small eyes and cardiac edema (**Figure 4B**), similar to published *foxc1a* mutant phenotypes (Ferre-Fernandez et al., 2020). We have previously shown that *foxc1a^p162^* mutant embryos exhibit early defects in forming the first aortic arch leading to a loss of circulation (Shin *et al*., 2019). Accordingly, we did not observe circulation in any *foxc1a^p162/Δ^* embryos, while heterozygous and homozygous wild type siblings were relatively normal (**Figure 4C**). The low penetrance of circulatory defects in homozygous and heterozygous wild type embryos suggests mild toxicity from Cre itself in these experiments. As with the *gata2a^fl^* allele, *foxc1a^fl/fl^* homozygous individuals appeared normal and could be grown to adulthood. Furthermore, *foxc1a^fl/fl^*embryos injected with zf-*cre* mRNA also exhibited loss of aortic arch and trunk circulation, while similarly injected *foxc1a^fl/+^* siblings were largely normal (**Supplementary Movies S3, S4**). Consistent with targeted knockout of *foxc1a*, we observe a nearly complete loss of *foxc1a* transcript expression following injection of *foxc1a^fl/fl^* larvae with *zf-cre* mRNA, while expression of the *foxc1b* paralog is unchanged (**Figure 4D**). Together, these observations demonstrate that *foxc1a^fl^* behaves as a conditional loss-of-function allele following Cre-mediated deletion.

**Figure 4.**
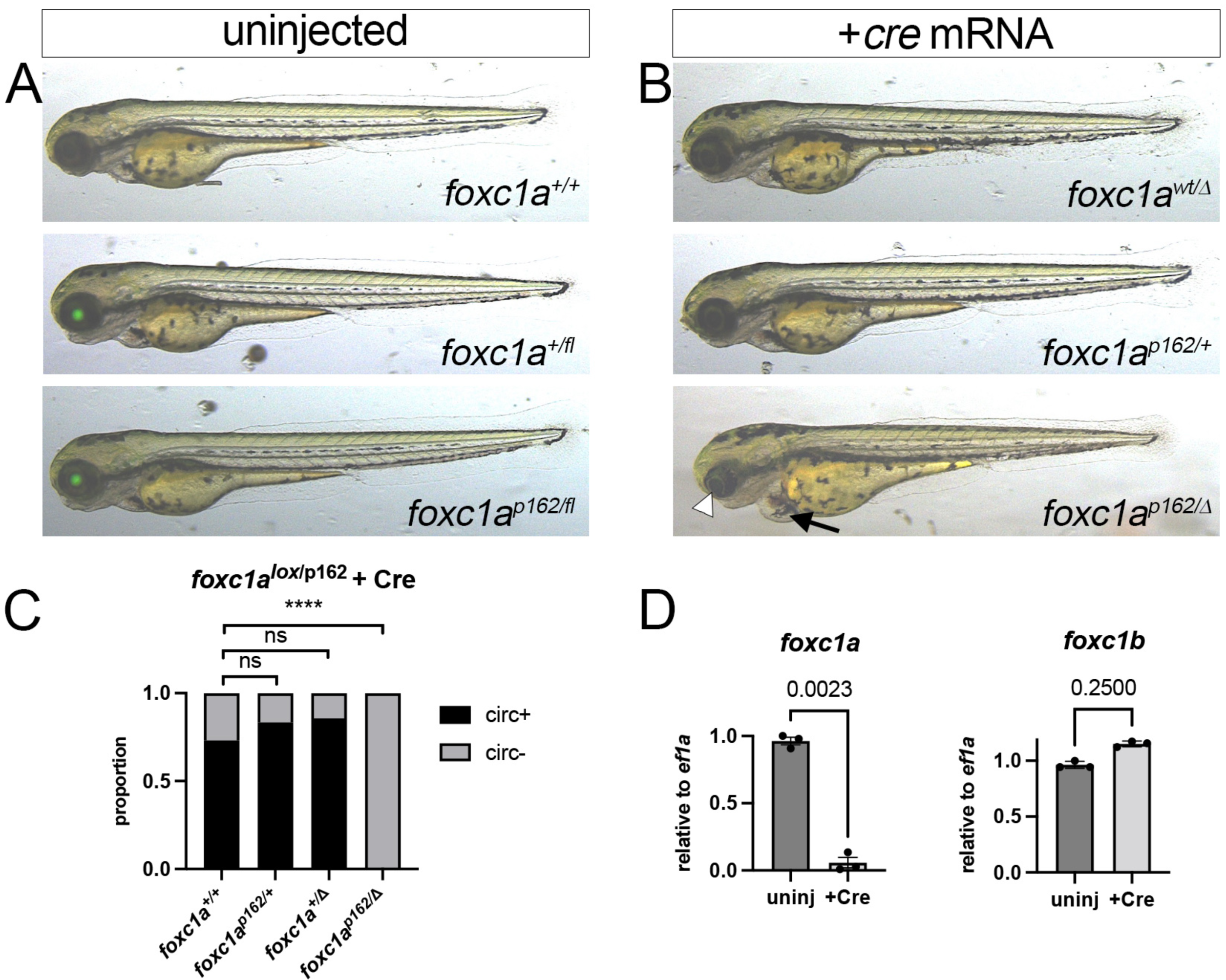
Functional validation of *foxc1a^fl^*. (A, B) Overlay of transmitted light and green fluorescence at 3 dpf. Larvae from cross between *foxc1a^+/fl^*and *foxc1a^+/p162^* carriers. (A) Uninjected embryos of indicated genotype with normal morphology. (B) Embryos injected with *cre* mRNA. Arrowhead denotes small eye and arrow indicates cardiac edema in bottom panel. (C) Proportion of 3 dpf larvae of indicated genotype with or without circulation. Total embryos: *foxc1a^+/+^* - 26, *foxc1^p162/+^* - 12, *foxc1a^+/Δ^* - 14, *foxc1a^p162/Δ^* - 16. Fisher’s exact test, ****p<0.0001. (D) Quantitative RT-PCR for *foxc1a* and *foxc1b* in *foxc1a^fl/fl^* embryos. Embryos were left uninjected or were injected with *zf-cre* mRNA at 1-cell stage. Paired t-test, p values are shown.

### Endothelial cell-specific *gata2a* knockout leads to defective lymphatic valve development

Floxed alleles allow for cell type- and/or stage-specific gene knockout. To demonstrate this utility in zebrafish, we used the *gata2a^fl^*allele to determine the endothelial autonomous requirement for *gata2a* during development. For this purpose, we first developed a line in which inducible Cre recombinase (CreERT2, referred to as “iCre”) was expressed with an endothelial-specific regulatory element. We have shown that 6 copies of the core conserved sequence from an intronic *gata2a* enhancer upstream of a basal promoter (referred to as *gata2aECE*) drives expression specifically in endothelial cells at early stages and becomes restricted to facial lymphatic endothelial cells by 6 dpf (Shin *et al*., 2019). Therefore, we used *gata2aECE* to establish a transgenic line expressing iCre, along with a *cryaa:mcherry* cassette to identify carriers (**Figure 5A**). We identified a founder from which progeny embryos displayed endothelial-specific recombination (*Tg(gata2aECE:CreERT;cryaa:mcherry)^um337^*; referred to as *gata2aECE:iCre*) and bred this line out to obtain a single copy insertion (**Figure 5B**). To confirm inducible endothelial-specific Cre activity, we crossed *gata2aECE:iCre* to fish bearing *Tg(ubb:loxP-cerulean-loxP;h2b-cherry)^jh63^* (Zhang et al., 2017), in which cells expressing Cre will switch on expression of a nuclear-localized form of mCherry (referred to as *ubb:SwitchRed*). Double transgenic *gata2aECE:iCre;ubb:SwitchRed* embryos show endothelial-specific H2B-mCherry expression at 6 dpf when exposed to 5 μM 4-hydroxytamoxifen (4OHT) from 1 to 3 dpf (**Figure 5C, D**). We observed H2B-mCherry in endothelial cells lining the facial lymphatic vessels, as well as those in aortic arch and brain blood vessels (**Figure 5C, D**). In trunk vessels, H2B-mCherry appeared to be restricted to endothelial cells within the posterior cardinal vein and the thoracic duct, with mosaic expression in intersegmental vessels and very few cells apparent in the dorsal aorta (**Figure 5D**). In *gata2aECE:iCre;ubb:SwitchRed* embryos treated with 4OHT from 2 to 3 dpf, H2B-mCherry was expressed in most endothelial cells within the facial lymphatic vessels, but expression in brain and aortic arch vessels was highly mosaic (**Figure 5E**). H2B-mCherry was similarly mosaic in posterior cardinal vein and thoracic duct endothelial cells but was not detectable in other blood vessels in the trunk (**Figure 5F**). Thus, *gata2aECE:iCre* drives endothelial-specific recombination that is progressively restricted to facial lymphatic endothelial cells as development proceeds.

**Figure 5.**
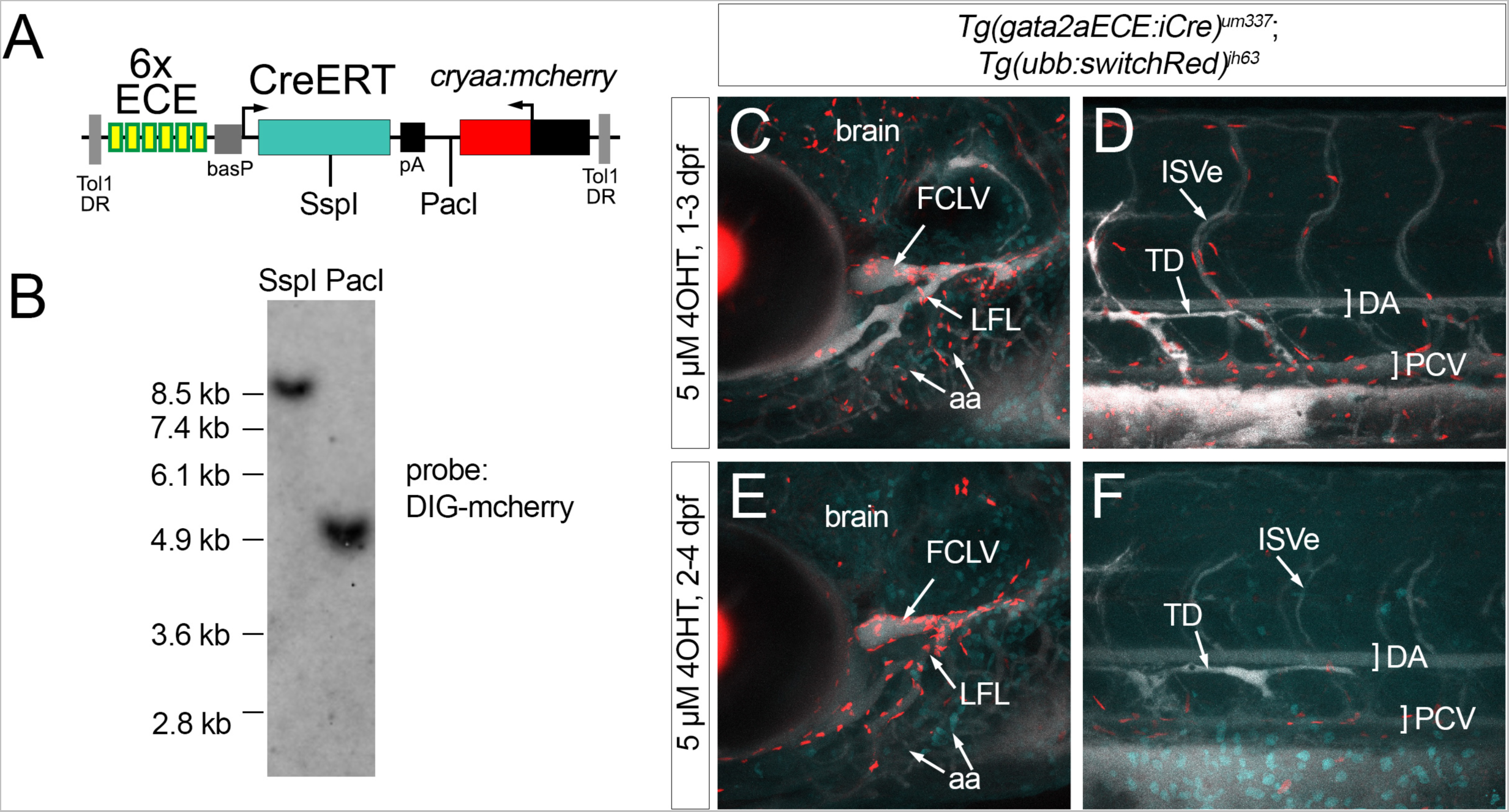
Generation of an endothelial-specific inducible Cre transgenic line. (A) Tol1 *gata2aECE:iCre* transgene. Locations of restriction enzyme sites are indicated. (B) Southern analysis using genomic DNA from *Tg(gata2aECE:iCre)^um337^* embryos. Blot was hybridized with a DIG-labeled *mcherry* probe and detected by chemiluminescence. (C-F) Confocal micrograph overlays of *Tg(gata2aECE:iCre)^um337^; (ubb:switchRed)^jh63^* larvae at 7 dpf showing expression of H2B-Cherry (red), cerulean (blue) and Qtracker705 introduced via lymphangiography (white). Lateral views, anterior to the left, dorsal is up. (C, E) Cranial vessels; (D, F) trunk vessels. FCLV – facial collecting lymphatic vessel, LFL – lateral facial lymphatic vessel, aa – aortic arch blood vessels, ISVe – intersomitive vein, TD – thoracic duct, DA – dorsal aorta, PCV – posterior cardinal vein.

To assess the requirement for *gata2a* in endothelial cells, we incrossed *gata2a^+/fl^;gata2aECE:iCre* adults and treated resulting progeny with DMSO or 4OHT from 1 to 3 dpf. Homozygous *gata2a^fl/fl^*larvae without iCre, identified by *cryaa:egfp* but not *cryaa:mcherry* and confirmed with genotyping, exhibited inflated swim bladders and normal circulation (**Figure 6A, C**). By contrast, 4OHT-treated *gata2a^fl/fl^*;*gata2aECE:iCre* larvae (referred to hereafter as *gata2a^iΔEC^*) displayed edema around the eyes and gut at a penetrance similar to that we previously reported for *gata2a^um27^*(Shin *et al*., 2019)(**Figure 6B, D**). In these experiments, we initially identified *gata2a^iΔEC^* larvae by lens expression of mCherry and Egfp, followed by genotyping for *gata2a^fl/fl^*. In contrast to *cre* mRNA injection (see **Figure 2F**), *gata2a^iΔEC^*larvae retain *cryaa:egfp* due to the specificity of CreERT expression. Interestingly, 4OHT treatment from 2 to 4 dpf resulted in much lower penetrance of the edema phenotype (**Supplementary Figure S4**). Notably, *gata2a^iΔEC^* larvae were otherwise similar to *gata2a^fl/fl^* siblings, with inflated swim bladders and normal trunk circulation (**Figure 6A, B**), unlike *gata2a^um27^* mutants (Shin *et al*., 2019). Furthermore, *gata2a^iΔEC^* embryos treated from 6 hpf to 24 hpf with 4OHT exhibited normal *urp1* expression in ventral KA neurons (**Figure 6E, F**), unlike *gata2a^fl/fl^* embryos injected with *zf-cre* mRNA (**Figure 2I**). Quantification of *gata2a* levels in H2B-mcherry-positive cells isolated by fluorescence activated cell sorting (FACS) from *gata2a^iΔEC^* larvae bearing *ubb:SwitchRed* (i.e. cells having exhibited Cre-mediated recombination) showed significant reduction in transcript containing exon 5, but not the 3’ UTR, compared to wild type or *gata2a^+/iΔEC^*larvae (**Figure 6G**). In contrast to *gata2a^fl/fl^*embryos injected with *zf-cre* mRNA, wild type transcript was still detectable in *gata2a^iΔEC^* larvae suggesting that recombination was robust, but incomplete when using *gata2aECE:iCre* (compare with **Figures 2K and 2G**).

**Figure 6.**
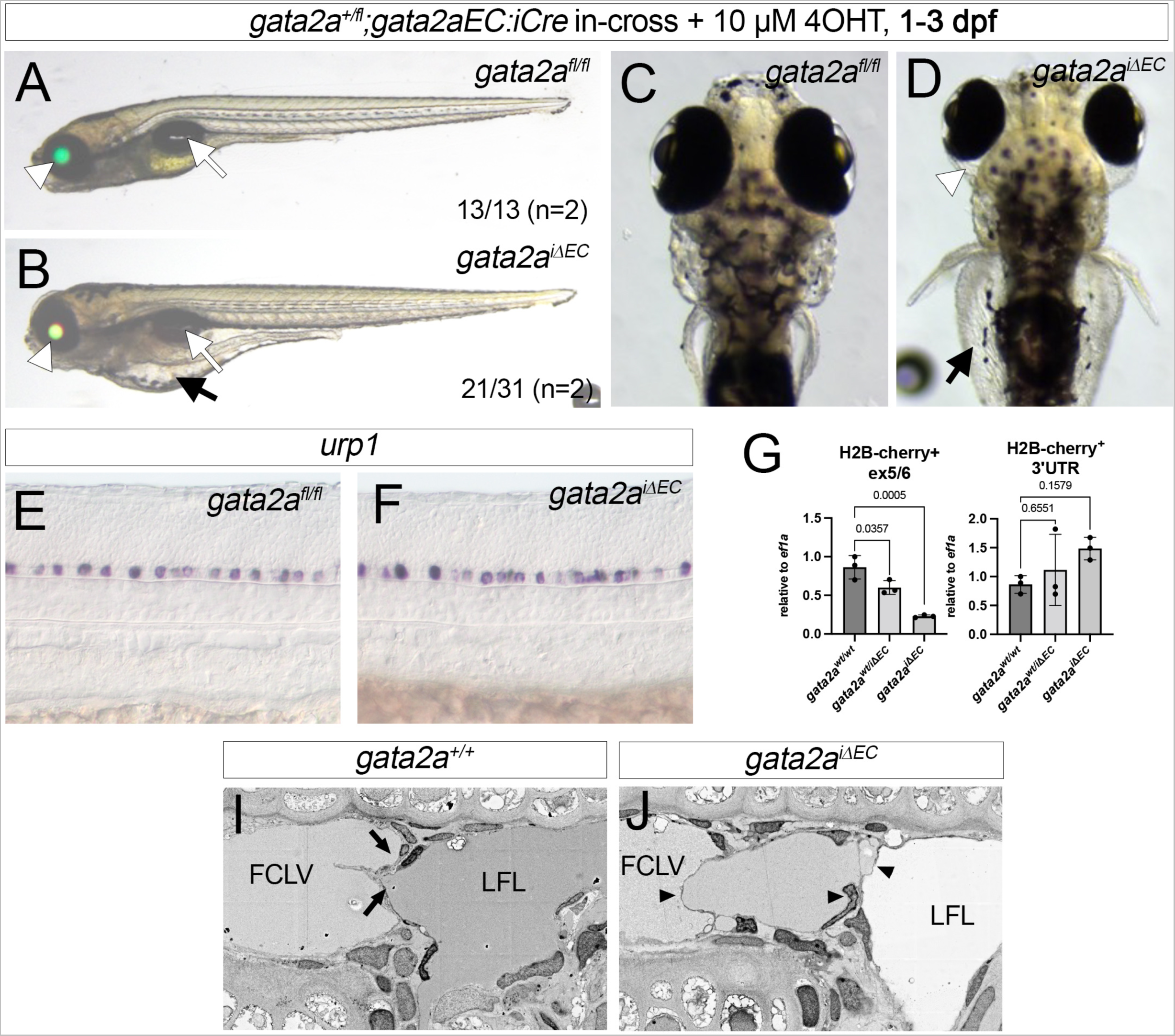
Endothelial *gata2a* is required for lymphatic valve function and formation. (A, B) 6 dpf and (C, D) 7 dpf larvae from indicated cross and 4OHT treatment at 10µM from 1 dpf to 3 dpf. (A, C) *gata2a^fl/fl^* larvae without iCre. (B, D) *gata2a^fl/fl^; Tg(gata2aEC:iCre)^um337^* larvae, referred to as *gata2a^iΔEC^*. (A, B) Overlay of transmitted light, green and red fluorescence. White arrowhead denotes lens, white arrow shows inflated swim bladder. Black arrows in (B, D) indicate edema around the gut. White arrowhead in (D) indicates edema around the eye. (E, F) *Urp1* expression by whole mount *in situ* hybridization in embryos of indicated genotype at 24 hpf. Embryos were treated with 4OHT at 5 µM from 6 hpf. (G) Quantitative RT-PCR for *gata2a* exon 5/6 or 3’ UTR from H2B-mcherry-positive cells isolated by FACS from *gata2a^iΔEC^;ubi:switchRed* at 5 dpf and 6 dpf following 4OHT treatment at 5 µM from 6 hpf to 4 dpf. For exon 5/6, analysis of variance, p<0.005; for 3’, not significant. Pairwise comparisons performed using Dunnett’s multiple comparison test, p values are indicated. (I, J) Scanning electron micrographs of lymphatic valves in embryos of indicated genotype at 7 dpf following 4OHT treatment at 5 µM from 1 dpf to 3 dpf. Lateral views, dorsal is up, anterior to the left. Both leaflets in normal bicuspid valve are indicated by black arrows. Multiple abnormal leaflets are indicated by black arrowheads in *gata2a^iΔEC^* larvae. FCLV – facial collecting lymphatic vessel. LFL – lateral facial lymphatic vessel.

We have previously shown that edema in *gata2a^um27^* mutant embryos coincides with defects in lymphatic valve formation (Shin *et al*., 2019). Therefore, we performed electron microscopy to investigate lymphatic valve morphology. In wild type larvae at 5 dpf, we observed the presence of flattened bicuspid valve leaflets separating the facial collecting lymphatic vessel (FCLV) and lateral facial lymphatic vessel (LFL), as previously described (**Figure 6I**, (Shin *et al*., 2019). By contrast, lymphatic valve leaflets appeared disorganized in *gata2a^iΔEC^* larvae that had been treated with 4OHT from 1 to 3 dpf. We observed multiple leaflets spanning the lumen between the facial collecting lymphatic vessel (FCLV) and lymphatic facial lymphatic (LFL) in a disorganized manner (**Figure 6K**), similar to what we have previously observed in *gata2a^um27^* mutant larvae (Shin *et al*., 2019).

## DISCUSSION

In this work, we identified points of improvement in existing zebrafish knock-in protocols to facilitate creation of floxed alleles in a single generation. We found that two particular steps were important. First, simultaneous CRISPR targeting of two genomic sites promoted complete replacement of endogenous sequence with a floxed version. Previous studies have relied on a single DSB to insert a reporter transgene, or splice trap cassette (Hoshijima *et al*., 2016; Li et al., 2019; Sugimoto et al., 2017; Wierson et al., 2020). However, a floxed allele requires an insertion at two distinct locations within a target locus. Furthermore, the targeting vector design in this case will bear homologous sequence (e.g. the target exon) between the loxP sites, providing a potential template for HDR. Indeed, at the *gata2a* locus a single DSB resulted in only partial knock-in, likely due to a short internal homology stretch (340 bp; **Figure 1B**) between the 5’ loxP and CRISPR sites. We reasoned that simultaneous DSBs at both loxP insertion points in the target locus would force HDR to use homology sequence outside of the target construct loxP sites. This worked successfully at the *foxc1a locus* to generate a floxed allele in a single generation. It is not possible to determine if knock-in occurred as a single repair event in our study since the internal *foxc1a* sequence was completely matched to the haplotype of injected embryos. However, given the low rate of targeted insertion observed with a single DSB in *gata2a*, the likelihood of two independent HDR mediated repair events occurring in *cis* would be exceedingly rare.

Based on the prevalence of off-target integrations we observed, a second improvement we made was to include a *myl7:mcherry* cassette outside of the homology arms as a screenable marker for these events. Unfortunately, this strategy did not work reliably in injected embryos, likely due to the high rate of multicopy transgene insertion in this setting. Indeed, *gata2a^um295^* and *foxc1a^fl^*founders carried both on- and off-target integrations, indicating that multiple independent germline insertions can occur in a single individual. Thus, the predominance of random insertions expressing both *cryaa:egfp* and *myl7:mcherry* over HDR-mediated knock-in events likely masks identification of the latter following injection. By contrast, counter-screening of P0 progeny identified individuals with off-target integrations, although it was not definitive: two putative *foxc1a^fl^*founders gave *cryaa:egfp-positive*/*myl7:mcherry*-negative larvae, but only one proved positive for the knock-in allele. Therefore, we would strongly recommend counter-screening along with 5’ and 3’ junction PCR, and Southern blot analysis to definitively identify carriers of a single copy floxed allele that do not have off-target integrations.

In our targeting construct, we matched homologous sequences with the haplotype of injected embryos, as previously recommended (Hoshijima *et al*., 2016). Sequencing the target locus should be the first step to ensure proper design of PCR primers, CRISPR targeting sites, and homology arms, all of which could be affected by sequence variants. While we did not directly test the effects of variants, there is considerable variability between and within wild type zebrafish strains (Balik-Meisner et al., 2018; Butler et al., 2015) and minor sequence variants in mouse can significantly reduce knock-in rates (Deng and Capecchi, 1992). In contrast to other studies, we also employed Cas12a instead of Cas9 for introducing DSBs. Unlike Cas9, which yields blunt ends, Cas12a leaves overhangs, while its PAM sequence lies opposite its cleavage site and the spacer can be up to 23 nucleotides (Zetsche et al., 2015). Thus, initial indels created by Cas12a-stimulated cleavage can often be re-targeted. Together, these characteristics likely contribute to the ability of Cas12a to stimulate higher HDR rates than Cas9 in both zebrafish and mammalian cells (Moreno-Mateos *et al*., 2017; Shahbazi *et al*., 2019). The TTTN PAM used by Cas12a also favors its application for targeting AT-rich non-coding sequences used for intronic loxP insertion sites in floxed alleles.

Our results using *gata2a^fl/fl^* underscore the utility of floxed alleles for genetic analysis at zebrafish larval stages. We previously relied on the *gata2a^um27^* allele, which causes numerous embryonic defects that can confound analysis at larval stages (Shin *et al*., 2019; Zhu *et al*., 2011), a common issue with most zebrafish mutants. Notably, *gata2a^um27^*mutant embryos have circulatory defects that could indirectly affect lymphatic valve development, which is sensitive to mechanosensory input in other models (Sabine et al., 2012). By contrast, *gata2a^iΔEC^* larvae displayed normal circulation and neuronal specification, yet exhibited defects in lymphatic valve function and morphogenesis with a severity and penetrance similar to *gata2a^um27^* (Shin *et al*., 2019). This observation is consistent with the known endothelial autonomous role for mouse *Gata2* in lymphatic valve formation (Kazenwadel et al., 2015). Of note, normal circulatory function in *gata2a^iΔEC^* embryos was somewhat surprising given the severity of circulation defects in *gata2a^um27^*embryos and endothelial expression of *gata2a* in embryos (Shin *et al*., 2019; Zhu *et al*., 2011). It is possible that *gata2a* acts at a stage prior to the 4OHT treatment used in our experiments. Alternatively, *gata2a* may function only in particular endothelial subtypes not targeted by *gata2aECE:iCre*. For example, *gata2aECE:iCre* did not induce recombination in arterial endothelial cells, raising the possibility that only this cell type in the trunk blood vessels is affected by loss of *gata2a*. The possibility also remains that circulatory defects in *gata2a^um27^* embryos are not endothelial autonomous. In any case, the availability of the *gata2a^fl^*allele, together with other Cre lines, now allows us to better investigate the cellular nature of the early circulatory defects.

The development of efficient CRISPR nucleases now enables zebrafish researchers to engineer knock-in alleles. Initial efforts in this regard introduced short epitope tags or reporter transgenes using a single DSB to insert exogenous sequence (Hoshijima *et al*., 2016 Wierson, 2020 #964; Hruscha *et al*., 2013). More recent studies used this approach to generate conditional loss-of-function alleles (Li *et al*., 2019; Liu et al., 2022). Most efforts have used a single DSB with a targeting vector to insert a conditional splice trap cassette into an intron via HDR (Li *et al*., 2019; Liu *et al*., 2022; Sugimoto *et al*., 2017). These alleles also incorporate reporter transgenes to observe endogenous expression and to assess Cre-mediated recombination. However, splice traps may not always lead to loss-of-function and reliable reporter expression is limited to strongly expressed genes. Moreover, splice traps will not work for small genes comprising only one or two exons. In general, a floxed allele is more likely to be definitively loss-of-function. Although these alleles can be more challenging to make, the improvements we have identified should aid researchers in this regard. The mouse community currently benefits from thousands of floxed alleles and hundreds of Cre lines allowing genetic analysis in any setting. These tools have allowed the mouse to become the preeminent genetic model for studying vertebrate development. The zebrafish is currently constrained by the lack of such tools. We hope that our efforts here can serve as an initial framework to establish more floxed lines that would allow conditional gene manipulation in zebrafish.

## ACKNOWLEDGEMENTS

This work was supported by R35HL140017 and R21OD030004 to N. D. L. from the National Institutes of Health. K.L. and S.A.W. were supported in part by grants from the National Institutes of Health (R01GM115911, R01HL150669, and 5F31HL147482). We thank Sarah Oikemus for critical reading of the manuscript.

## METHODS

### Zebrafish

Zebrafish were handled according to approved University of Massachusetts Chan Medical School Institutional Animal Care and Use Committee protocols. Establishment and characterization of *gata2a^um27^*, *foxc1a^p162^*, and *Tg(ubi:CtoH2b-cherry)^jh63^* lines has been described elsewhere (Banerjee *et al*., 2015; Zhang *et al*., 2017; Zhu *et al*., 2011). Generation of all other alleles is described below.

### Target locus sequencing

We sequenced target loci in genomic DNA isolated from finclips from 12 individual TL wild type fish (6 males and 6 female). For *gata2a*, we amplified by PCR using superPfx (Cowin Biotech) using primers: 5’-GGGGAATTGCGCTGTACGCCTATAAA-3’ and 5’-AATATGGACCTTTGAGGTACCACCCC-3’ to obtain genomic sequence flanking exon 5 (GRCz11, chr11:3,849,652-3,852,157). For *foxc1a*, we used primers: 5’-CTTAACCTCGCTGTATGGCTGTAG-3’ and 5’-CCCACAAGGCAGTGGCTAGCAGAACTCATG-3’ to amplify a spanning genomic sequence (chr2:686,529-689,296) of upstream including 5’ homologous arm, promoter, coding exon and 3’UTR, and 5’-CGGGAATAACAGCTGTCAAATGTC-3’ and 5’-AACCGCTGTAATCACTACCAACCGCTCATG-3’ to amplify 3’ homologous sequence (chr2:685,341-687,455). Amplicons were cloned into pBluescript linearized with SmaI. These were used for further plasmid construction for making targeting vectors describing below. We sequenced individual clones from each ligation and aligned them to reference genomic sequence (GRCz11). For embryo generation, we used a male and a female bearing the same haplotype for each locus that also matched the homology arms in the respective targeting construct (see Supplementary Files 1 and 2).

### Plasmid construction

For generation of a *gata2a* targeting vector, 5’ and 3’ homologous arms corresponding to GRCz11 chr11:3,851,035-3,852,157 and chr11:3,849,652-3,850,708, respectively, were PCR-amplified from sequence validated pBluescript clones described above. These PCR products were combined with a gBlock encoding *gata2a* genomic sequence comprising exon 5 (chr11:3,850,717-3,851,034), a 5’ loxP sequence (IDT), FRT-flanked cryaa:egfp, and a 3’ loxP site in a HiFi Assembly reaction (NEB) with pBS-ISceI to give pBSIce-gata2aKI (**Supplementary File 1**).To construct a *foxc1a* targeting vector, 5’ loxP sequence and point mutations (5’-TTTC-3’ to 5’-CCCT-3’) on the Cas12a RNP1 PAM sequence were added in the upstream of the promoter sequence (chr2:689,007) in pBluescript vector described above. DNA fragments of modified *foxc1a* locus with 5’ homologous arm (chr2:689,008-689,404) and loxP, and 3’ homologous arm (chr2:685,632-686,723) were separately PCR-amplified from the vectors. After digestion of pKHR8 (Addgene#74625; Hoshijima et al. 2016) using BamHI and SalI in MCSs, the DNA fragments were assembled using NEbuilder HiFi DNA assembly kit to give pKHR8-foxc1aKI (**Supplementary File 2**). A Cre Gateway middle entry clone was generated by PCR using 5’-GGGGACAAGTTTGTACAAAAAAGCAGGCTTGACCaTGCCCAAGAAGAAGAGG-3’ and 5’-GGGGACCACTTTGTACAAGAAAGCTGGGTAatcgccatcttccagcaggcg-3’, followed by BP cloning into pDONR221 (ThermoFisher), to give pME-CreNS. pCS-CreNS was constructed via a multisite LR reaction using pCSDest2, pME-CreNS and p3E-MCS1 (Moore et al., 2013; Villefranc et al., 2007). Alternatively, we used a Cre with zebrafish optimized codon usage (zfCre). For this purpose, we first constructed pME-zfCreERT^2^ by HiFi assembly using two gBlock DNA fragments (IDT) and pME-MCS1 vector (Kwan et al., 2007) linearized by inverse PCR using primers (5’-GTGAGTCGTATTACATGGTCATAGCTG -3’ and 5’-GGCCGTCGTTTTACAACGTCGTGACTG -3’). To generate pME-zfCre, pME-zfCreERT^2^ was digested using XhoI to remove zfERT^2^ and self-ligated. We constructed pCS-zfCre using pCSDest2, pmE-zfCre and p3E-MCS1 in a multisite reaction using LR Clonase II (Thermofisher). For transgenic expression of CreERT^2^, we first modified the Tol1 backbone vector, pToneDest (Addgene#67969)(Shin et al., 2016), by adding a *cryaa:mcherry* cassette to give pToneDestCryR. We then performed a multisite reaction using LR Clonase II with pToneDestCryR, p5E-6xgata2aECEbas (Shin *et al*., 2019), pENTR/D-CreERT2 (Addgene#27231)(Mosimann et al., 2011), and p3E-MCS1 to give pTol1-gata2aECE:CreERT^2^;cryaa:mcherry.

### *In vitro* transcription

To synthesize *cre* and *zf-cre* mRNA, we performed *in vitro* transcription using a mMESSAGEmMACHINE SP6 kit (ThermoFisher) with respective pCS vectors linearized with NotI (NEB). To synthesize Cas9 sgRNA, we annealed the following oligonucleotides: 5’-tagGAGAGGGACGAGCGAGGCC, 5’-aaaCGGCCTCGCTCGTCCCTCT and cloned them into pDR274 digested with BsaI to give pDR274-gata2a_3pex5sgRNA1. The resulting plasmids was linearized with HindIII (NEB) and used to synthesize sgRNA using the T7 MEGAscript kit (ThermoFisher). Preparation of DNA templates and generation of crRNAs targeting *gata2a* (5’ site for loxP insertion RNP target; 5’-ACATGACCATGGGGTTGTTCCTT-3’) and *foxc1a* (RNP1 target: 5’-CGATGCGCGCTCCGAGAGAAAGAG-3’; RNP2 target: 5’-CTGCGGCACACTTGAACGATCGTC-3’) with a full length direct-repeat for *Lachnospiraceae bacterium* Cas12a were performed as described previously (Liu et al., 2019), using bottom strand oligos (5’-TGACCATGGGGTTGTTCCTTATA ATCTACACTTAGTAG-3’, 5’-CTCTTTCTCTCGGAGCGCGCATCG ATCTACACTTAGTAG-3’ and 5’-GACGATCGTTCAAGTGTGCCGCAGATCTACACTTAGTAG-3’, respectively). Tol1 transposase mRNA was synthesized from linearized pToneTP as described elsewhere (Shin *et al*., 2016).

### SpCas9 and LbCas12a Protein purification

3xNLS-SpCas9 protein expression and purification was performed as previously described {Wu, 2019 #966}. LbCas12a-2xNLS protein expression and purification was performed as previously described (Liu et al., 2019).

### Line generation

For targeting *gata2a*, 50pg of pBSIce-gata2aKI, 60pg of sgRNA, 0.8pg of 3xNLS-SpCas9 protein and 1mU of I-SceI (NEB) was injected per embryo at early 1-cell stage. At 2 dpf, embryos were separated based on expression of EGFP in the lens. Genomic DNA from individual embryos was purified by adding 50µL of 0.05M NaOH, incubating at 98°C for 10 minute, and then adding 10µL of 0.5M Tris pH7.5 (HCl). Precise knock-in of 3’ homologous arm was evaluated by PCR using KAPA2G Fast HotStart ReadyMix (KAPA Biosystems) with a primer set (5’-CCACTAGTTCTAGAGCGGC-3’ and 5’-GAACTGTTGGCTACTAACACTAATACTG-3’) across the junction with endogenous sequence using genomic DNAs of individual embryos as templates. Subsequently, injected embryos with EGFP lens expression were grown to adulthood. Founders were screened by individual outcross with wild type adults followed by screening for lens expression of EGFP in progeny. The same primer set as above was used for PCR screening of genomic DNA from pooled embryos with lens EGFP expression. To delete the off-target loxP-flanked exon5 sequence, we injected 10 pg of *cre* mRNA into embryos from an incross of *gata2a^um295/um295^* parents and grew these to adulthood to give rise to *gata2a^um329^*. To insert a loxP site 5’ of the targeted *gata2a* exon 5 in *gata2a^um329^* fish, we designed oligodeoxynucleotides (ODN) to include loxP sequence flanked by XhoI and EcoRI restriction sites and 30bp of homology (IDT). ODNs were annealed in 50mM NaCl solution to generate double stranded ODN template (dsODN) by incubating at 98°C for 10 minutes followed by 95°C, 90°C, 85°C, 80°C, 75°C, 70°C, 65°C and 63°C each for 20 seconds before 62°C. We co-injected 45pg of dsODN and 57 fmol of LbCas12a-2xNLS RNP (see above) per embryo at early 1-cell stage. Embryos were derived from an incross of homozygous *gata2a^um329/um329^* adults. After injection, integration was evaluated by PCR using 5 PRIME HotMasterTaq DNA polymerase (Quantabio; primers : 5’-TTAGAAGAACCGGTCTTCAGTATG-3’ and 5’-CTGTAGGCAAGAAAGCAGAGTATTGTCA-3’) on genomic DNA of pooled or individual embryos followed by digest with XhoI (NEB). Following validation of knock-in, injections were repeated and embryos grown to adulthood. Founders were identified using the same PCR primers and XhoI digestion as above. For generation of *foxc1a^um383^*, 40 pg of pKHR8-foxc1aKI, 9.5 fmol of Cas12aRNP1, 9.5 fmol of Cas12aRNP2 and 1mU I-SceI were co-injected into wild type TL embryos at early 1-cell stage. Injected embryos were sorted at 2 dpf for expression of Venus in the lens and/or mCherry in the heart. Integration was evaluated by PCR using KAPA2G Fast HotStart ReadyMix on genomic DNA from individual embryos with primer sets that span 5’ and 3’ junctions between homology arms and endogenous sequence (5’ junction primers; 5’-CAGATTTTACCTCTGGGTATTATACGA-3’ and 5’-TACTGGCCACCTCTTATAACTTC-3’, 3’ junction primers; 5’-GATGAAGCTACATGGCTGTAGAACGTCAG-3’ and 5’-AACCGCTGTAATCACTACCAACCGCTCATG-3’). After initial analysis, injections were repeated and only embryos expressing both Venus and mCherry were grown to adulthood. Founders were screened by individual outcross to wild type followed by screening for Venus and mCherry expression at 3 dpf. Embryos were separated based on Venus and mCherry expression, then pooled and used to isolate genomic DNA. We performed PCR across 5’ and 3’ junction sequences as above. To generate an endothelial-specific inducible Cre line, we co-injected 25 pg of pTol1-gata2aECE:CreERT^2^ and 50 pg of Tol1 transposase mRNA into wild type 1-cell stage embryos. Only embryos expressing mCherry in the lens were grown to adulthood. Founders were identified by individual outcross and screening of progeny for mCherry lens expression at 3 dpf. Founders were subsequently validated by crossing to an adult bearing *Tg(ubi:CtoH2b-cherry)^jh63^* followed by tamoxifen treatment. A founder was identified that gave robust endothelial recombination and designated as *Tg(gata2aECE:CreERT^2^;cryaa:mcherry)^um337^*. Following successive outcrosses with wild type, *Tg(gata2aECE:CreERT^2^;cryaa:mcherry)^um337^* a single copy insertion of the transgene was confirmed by Southern blotting (see below).

### mRNA Injections

To delete loxP-flanked sequences we injected 10 to 25 pg of *cre* mRNA, or 6.25pg of *zf-cre* mRNA into incrosses of fish that were heterozygous or homozygous for *gata2a^um345^*or *foxc1a^um383^* as indicated in the text.

### Tamoxifen administration

A 10 mM 4-hydroxytamoxifen (4OHT; MilliporeSigma) stock solution was made by dissolving in DMSO, aliquoted, and stored at −80°C. For treatment, embryos were treated for the indicated timepoints by dissolving 10 mM stock to a final of 5 or 10 µM in standard egg water. In cases where larvae were to be subjected to confocal imaging, embryos were also treated with 0.003% phenylthiourea (PTU). Embryos were placed in eggwater with fresh 4OHT every 24 hours. After the indicated treatment period, embryos were washed several times and maintained in PTU until 6dpf for imaging. For scanning electron microscopy (SEM), embryos were treated with tamoxifen at 5 µM between 1 dpf and 3 dpf grown until 7 dpf. Larvae were then analyzed by SEM and genotyped as described previously (Shin *et al*., 2019).

### Fluorescent-activating cell sort (FACS)

FACS was performed at room temperature using cells dissociated from embryos carrying the *gata2a^um345/um345^*, *gata2a^+/um345^* or *gata2a^+/+^*, carrying transgene of the *Tg(gata2aECE:CreERT^2^;cryaa:mcherry)^um337^* and *Tg(ubi:CtoH2b-cherry)^jh63^* at 5 dpf or 6 dpf with 4OHT administration between 6 hpf and 4 dpf under PTU treatment. Briefly, embryos (n=300 to 500) in 1x TrypLE (ThermoFIsher) at 28.5°C were minced with razor blade, dissociated within 15 minutes using P1000 pipetman (Gilson), and strained using 40µm pore-sized cell strainer (Falcon). Trypsin was inactivated by adding fetal bovine serum and cells rinsed with Leibovitz’s L-15 media without phenol red (ThermoFisher). Cells were spun down at 300g for 5 minutes at room temperature and resuspended with collection solution (L-15 media/10% embryo extract) for FACS. mCherry-positive and CFP-negative cells (150 x 10^3^ to 350 x10^3^ cells) were sorted by Flow Cytometry Core Lab at UMass Chan Medical School.

### PCR and genotyping

Larvae from crosses between *gata2a^+/um27^* and *gata2a^+/um345^*, with or without *cre* mRNA injection, individual embryos were separated by *cryaa:egfp* expression and subsequently genotyped for *gata2a^um27^* using a KASP assay as previously (Biosearch Technologies) (Shin *et al*., 2019). Excision of floxed exon 5 was determined by PCR using the following primers: 5’-TTAGAAGAACCGGTCTTCAGTATG-3’ and 5’-GATCGCAGCCAAGCTTAACATTAAA-3’. *Gata2a^iΔEC^*larvae and embryos were identified using a KASP probe. For embryos from *foxc1a^p162/um383^*complementation cross, we first sorted embryos *cryaa:venus* expression at 3 dpf. Individual embryos were then genotyped for *foxc1a^p162^* by KASP assay, as previously (Shin *et al*., 2019) and for deletion of *foxc1a* using PCR with 5’-CTTAACCTCGCTGTATGGCTGTAG-3’ and 5’-CCCACAAGGCAGTGGCTAGCAGAACTCATG-3’. Genotyping of *foxc1a^fl^* was done by KASP assay.

### RT-qPCR

Total RNA was purified from Cre-injected or uninjected embryos (5 to 10 per sample) with TRIzol reagent (ThermoFisher). For FACS-isolated *mcherry+* cells (1.50-3.5×10^5^) we used the Allprep DNA/RNA Micro kit (QIAGEN). cDNA was synthesized using 10ng of total RNA with SuperScript III First-Strand Synthesis SuperMix (ThermoFisher). For *foxc1a*, we removed residual genomic DNA before the reverse transcription reaction by treating with TURBO DNase (ThermoFisher). qPCR was performed with PowerSYBR Green PCR Master Mix (ThermoFisher) using ΔΔCT method on StepOne Plus Real-Time PCR machine (ThermoFisher). We used *eef1l1* for normalization. Primers were as follows: *eef1l1* (5’-GTCTGCCACTTCAGGATGTGTAC-3’ and 5’-GCATCTCAACAGACTTGACCTCAG-3’; (Shin *et al*., 2019)), *gata2a* ex5/ex6 (5’-CACCACACTCTGGAGACGCAATG-3’ and 5’-CCTGTTGCGTGTCTGAATACCATC-3’), *gata2a* 3’UTR (5’-CCCCTAACATGCTGGAATACAC-3’ and 5’-CACTTCTCTGCTTGTATGTCGTATC-3’), *foxc1a* (5’-CGGGAATAACAGCTGTCAAATGTC-3’ and 5’-GGTCAAAATTTGCTGCAGTCATACAC-3’) and *foxc1b* (5’-CGGTGACCGGAAGCAATAGCTGTC-3’ and 5’-GGGTCAGAACTTGCTGCAGTCGT-3’).

### Southern analysis

We purified genomic DNA from pooled embryos using the Blood & Cell Culture DNA Midi kit (QIAGEN). 5 or 10 µg of DNA was fully digested overnight with indicated restriction enzyme at the appropriate temperature. Digested samples were loaded onto a 0.7% agarose gel made with Tris acetate EDTA buffer alongside 100-200 ng of digoxigenin (DIG)-labeled DNA marker VII (MilliporeSigma) and run at 25V for 18 hours. After electrophoresis, the gel was sequentially incubated in 250 mM HCl for 25 minutes, 0.5M NaOH/1.5M NaCl for 45 minutes, and then 0.5M Tris pH7.5-HCl/1.5M NaCl for 30 minutes. Genomic DNA was then transferred from the gel to positively charged nylon membrane (MilliporeSigma) by standard capillary transfer overnight. Membrane was denatured on filter paper soaked in 0.4M NaOH for 5 minutes and UV-crosslinked at 420 x 10^2^ µj/cm^2^ in an XL-1000 UV CrossLinker (Spectronics Corp). Membranes were then prehybridized with DIG Easy Hyb solution (MilliporeSigma) in a HybriBag (Cosmo bio) at 42°C for 30 minutes with agitation. Prehybridization solution was replaced with fresh DIG Easy Hyb solution containing DIG labeled DNA probe at a final concentration between 5-10 ng/ml and incubated overnight at 42°C with agitation in a hybridization incubator (Robbins Scientific). An EGFP probe was PCR-amplified (5’-ATGGTGAGCAAGGGCGAGGAGCTG-3’ and 5’-ACTTGTACAGCTCGTCCATGCCG-3’) with DIG-labeled dNTP mix (MilliporeSigma) and AccuStart II DNA polymerase (Quantabio) using pBSIce-gata2aKI as a template followed by gel purification. Venus and mCherry fragments were PCR-amplified (*venus*:5’-ATGGTGAGCAAGGGCGAGGAGCTG-3’ and 5’-GGCGGCGGTCACGAACTCCAG-3’; *mcherry*: 5’-ATGGTGAGCAAGGGCGAGGAG-3’ and 5’-CTTACTTGTACAGCTCGTCCATG-3’) from pKHR8-foxc1aKI and separately subcloned into pJET2.1 (ThermoFisher) to give pJET-mVenus and -mCherry, respectively. DIG-labeled *venus* and *mcherry* DNA probes were then generated as above. Following hybridization, membranes were washed with 2x SSC/0.1% SDS at room temperature for 5 minutes, 0.5x SSC/0.1% SDS at 65°C for 20 minutes and 0.25x SSC/0.1% SDS at 65°C for 20 minutes, with agitation. After rinsing with maleic buffer (0.1M Maleic acid/150mM NaCl/0.3% Tween20-pH7.5), membrane was blocked in 1x Casein blocking buffer (MilliporeSigma) diluted in maleic buffer at room temperature for 30 minutes and incubated with anti-DIG antibody conjugated with alkaline phosphatase (1:50,000 dilution; MilliporeSigma) in blocking buffer at room temperature for 30 minutes. Membranes were washed with maleic buffer at room temperature for 15 minutes twice, preactivated with 0.1M Tris-pH9.5/0.1M NaCl/50mM MgCl_2_ then reacted with CDP-star ready-to-use substrate (MilliporeSigma). Chemiluminescence was detected by exposing membrane to film using CL-XPosure (ThermoFisher) or direct imaging on a ChemiDoc (Bio-rad).

### *In situ* hybridization

DNA templates for *urp1* or sst1.1. were generated by PCR as described elsewhere (Andrzejczuk *et al*., 2018). Following PCR, fragments were gel purified and then used as templates to generate DIG-labeled RNA probes using T3 polymerase. *In situ* hybridization was performed as previously described (Hauptmann and Gerster, 1994).

### Imaging

Confocal imaging was performed using a LMS 710NLO (Zeiss) with ZEN10. Prior to imaging, larvae were subjected to lymphangiography using Qtracker 705 quantum dots as described previously (Shin *et al*., 2019). Bright field and fluorescent images ware taken on M165FC (Leica) equipped with AxioCamMRc (Zeiss) using AxioVision SE64 Rel 4.9.1 software (Zeiss). Videos of blood circulation were acquired using differential interference contrast imaging on an Axioskop2 Plus (Zeiss) equipped with DMK21AU04 (IMAGINGSOURCE) using IC capture 2.5 software (IMAGINGSOURCE). Video files were saved as .mp4 and imported into Adobe Premiere Pro for labeling (Adobe). SEM sample preparation and image acquisition were performed as previously described (Shin *et al*., 2019).

### Statistical analyses

Statistical tests are noted in Figure legends.

## SUPPLEMENTARY FIGURES

**Figure S1.**
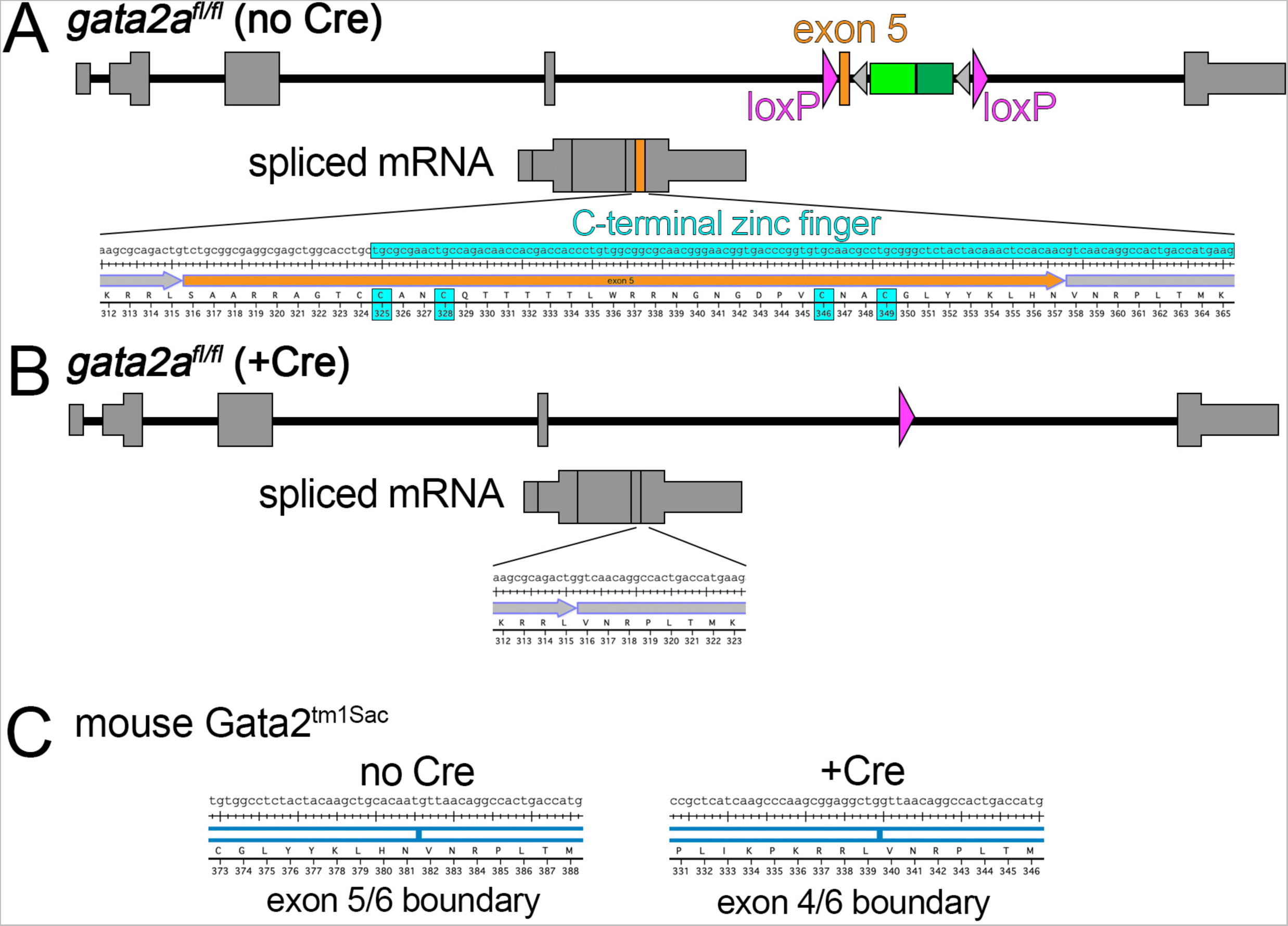
(A) Zebrafish *gata2a^fl/fl^* locus in the absence of Cre. Position of loxP sites flanking exon 5 are indicated. Structure of spliced mRNA is indicated with position of exon 5 denoted in orange. Sequence of exon 5 and flanking exon boundaries in *gata2a* transcript is shown. Coding sequence for C-terminal zinc finger is highlighted in cyan, as our individual cysteine residues within the zinc finger that are responsible for zinc binding. (B) Zebrafish *gata2a^fl/fl^* locus following Cre-mediated recombination. Spliced mRNA is indicated with highlight showing exon 4/6 boundary and in-frame coding sequence. (C) Sequence of indicated exon boundary in *Gata2* transcript expected from the mouse *Gata2^tm1Sac^* allele in the absence of presence or Cre.

**Figure S2.**
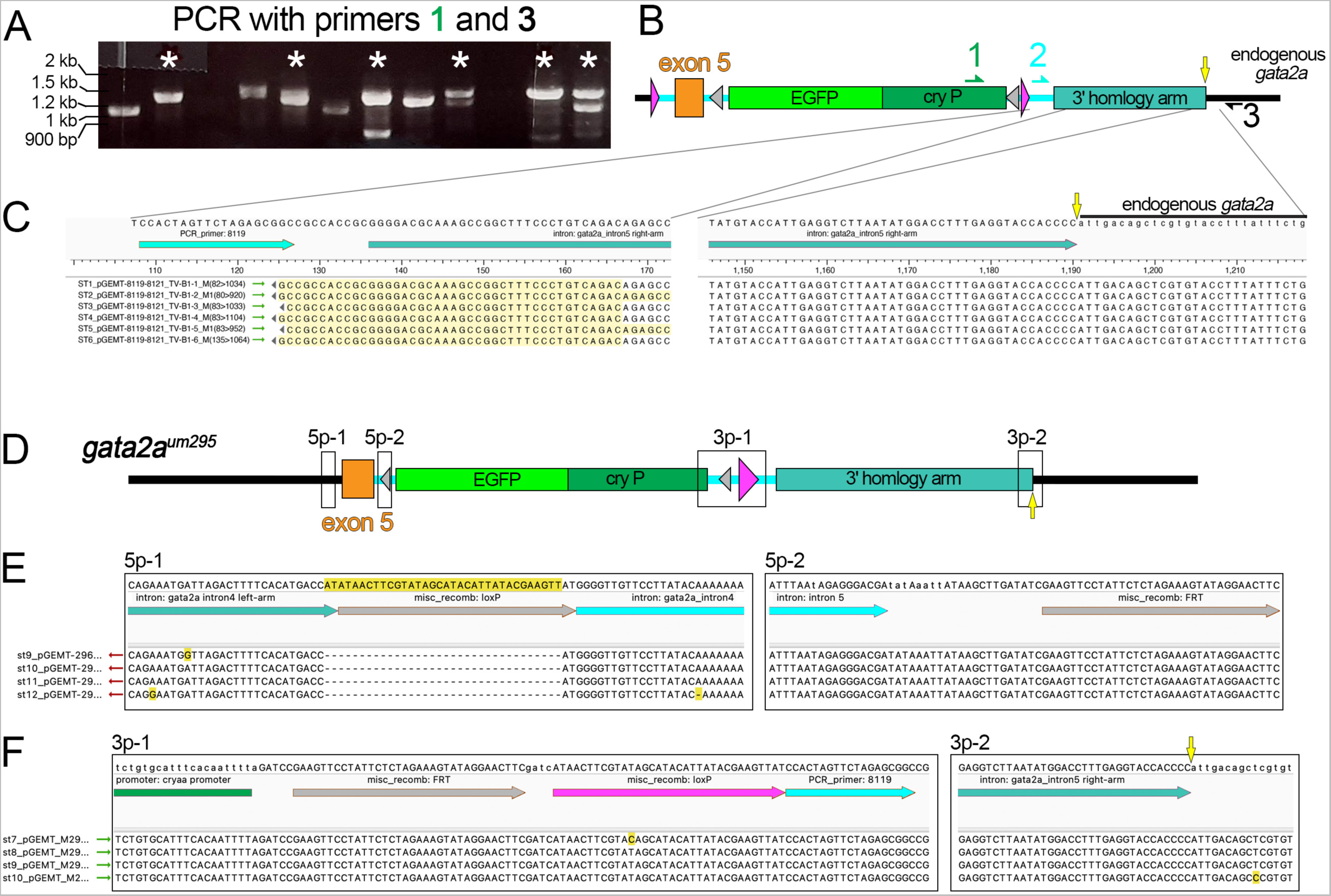
(A) PCR of individual *cryaa:egfp*-positive embryos injected with *gata2a* targeting construct, Cas9 RNP and ISce-I. Lanes with amplification of expected product size are indicated by an asterisk. PCR primers are indicated and their location is shown in (B). (B) Schematic of 3’ junction PCR at *gata2a* exon 5 target. (C) Alignment of cloned fragments spanning the 3’ homology arm and junction. Each sequence is a contiguous cloned fragment, of which only the 5’ and 3’ ends are shown. (B, C) Yellow arrow denotes junction between endogenous sequence and targeting construct homology arm sequence. (D) Exon 5 in *gata2a^um295^*. Labeled boxes denote regions for which sequence is shown in (E, F). (E). 5’ and 3’ sequence from cloned fragments spanning exon 5 aligned to *gata2a^fl/fl^*reference sequence. Note absence of 5’ loxP site from all 4 cloned fragments. (F) 5’ and 3’ sequence from fragments spanning the 3’ loxP site, across the homology arm sequence and into the endogenous *gata2a* locus. (D, F) Yellow arrow denotes junction between endogenous sequence and targeting construct homology arm sequence.

**Figure S3.**
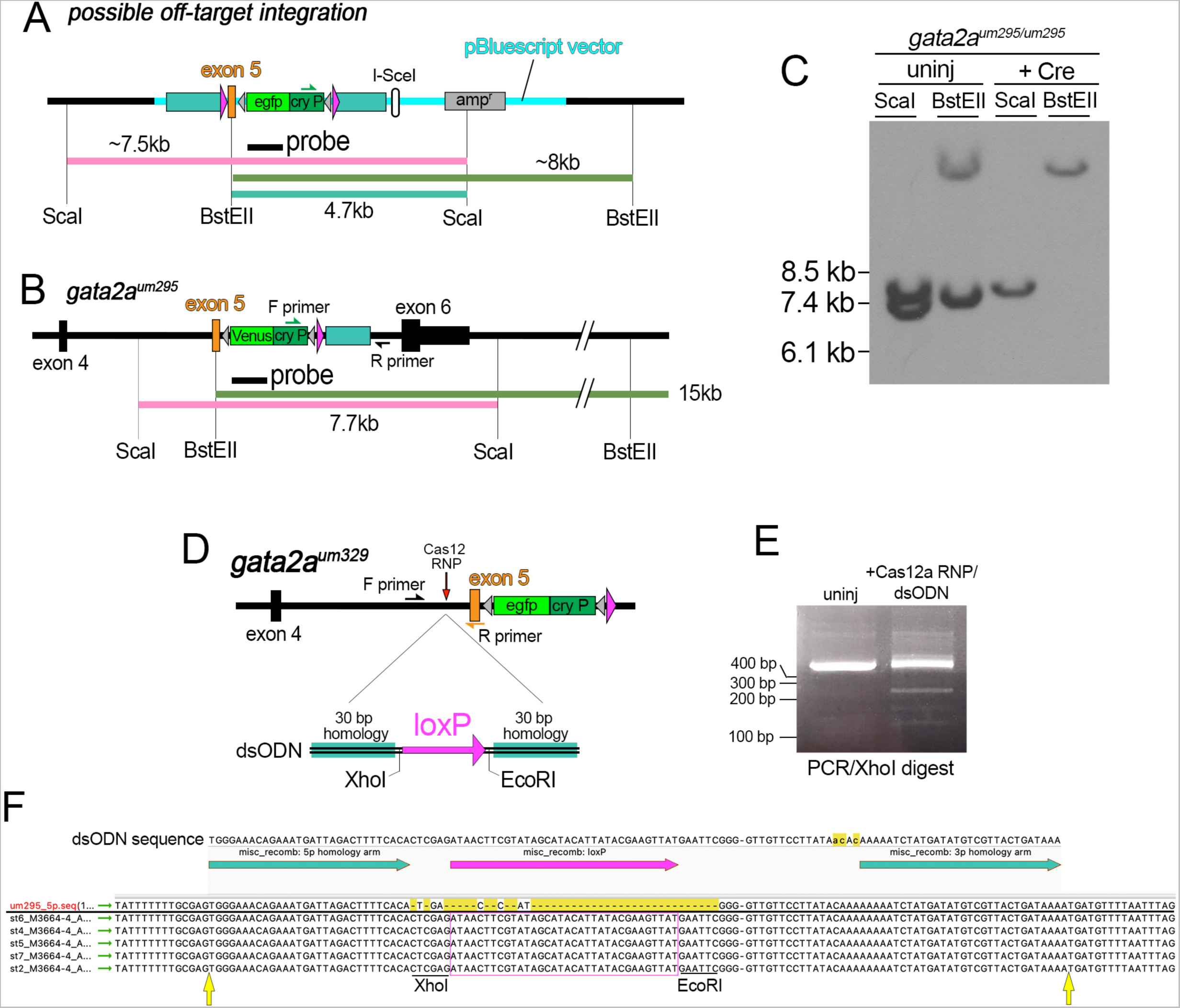
(A) Possible off-target integration arising from targeting vector cut only once by ISce-I following injection. (B) *gata2a^um295^* locus. (C) Southern blot with genomic DNA from homozygous *gata2a^um295^*embryos left uninjected or injected with *cre* mRNA. Blot was hybridized to a DIG-labeled probe for EGFP. (D) *gata2a^um329^* locus showing location of Cas12a RNP and dsODN structure used to insert the 5’ loxP site. (E) PCR product across insertion point for 5’ loxP site in embryos left uninjected or those injected with Cas12a RNP and loxP dsODN shown in (D). PCR products were digested with XhoI. Only products from embryos injected with Cas12a and dsODN show evidence of cutting, consistent with insertion of the exogenous sequence at the target site. (F) Sequence validation of the 5’ loxP insertion in *gata2a^fl/fl^*. Sequence of cloned fragments spanning the putative loxP insertion in F1 embryos from a P0 founder. Sequences are aligned to the *gata2a^um295^* as reference and the ODN sequence is shown. Yellow arrows denote junction points between ODN homology and endogenous sequences.

**Figure S4.**
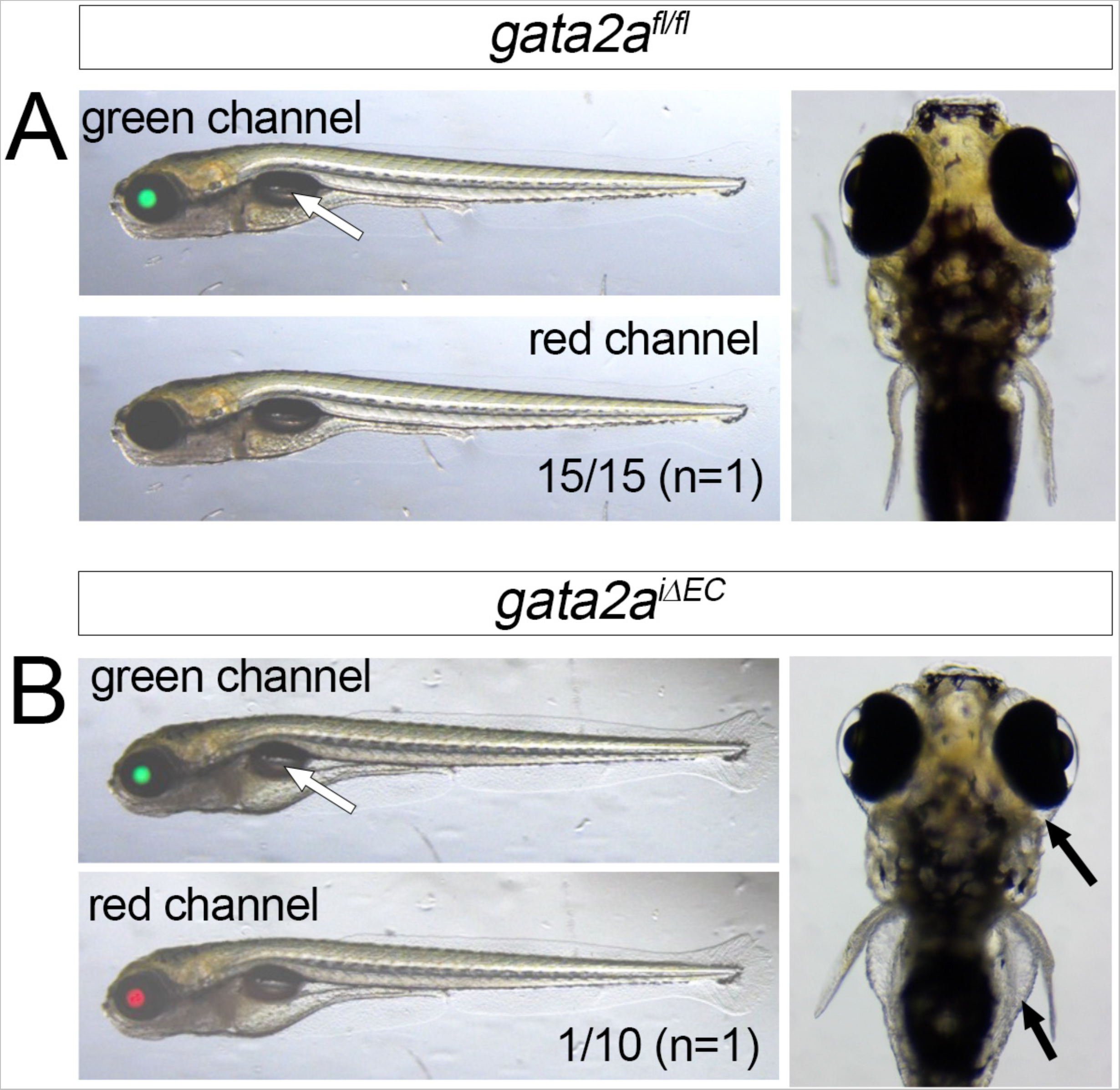
(A, B) Larvae at 6 dpf of indicated genotype. Left panels, overlay of transmitted light with indicated fluorescence channel. Lateral views, dorsal is up, anterior to the left. Right panels, dorsal view. White arrows indicate inflated swim bladder. (B) Arrows indicate mild edema around the gut and eye.

**Movie S1**. Heartbeat and circulation in a *gata2a^+/fl^*embryo at 2 dpf injected with *zf-cre* mRNA.

**Movie S2**. Heartbeat and circulation in a *gata2a^fl/fl^*embryo at 2 dpf injected with *zf-cre* mRNA.

**Movie S3**. Heartbeat and circulation in a *foxc1a^+/fl^* embryo at 2 dpf injected with *zf-cre* mRNA.

**Movie S4**. Heartbeat and lack of circulation in a *foxc1a^fl/fl^* embryo at 2 dpf injected with *zf-cre* mRNA.

**Supplementary Table 1.** Phenotypic scoring of embryos from cross between

| um27/exon5 genotype<sup>1</sup> |  | uninjected |  |  |  | +25 pg *cre* mRNA |  |  |  |
| --- | --- | --- | --- | --- | --- | --- | --- | --- | --- |
|  |  | cryG+ |  | cryG- |  | cryG+ |  | cryG- |  |
|  |  | sb+ | sb- | sb+ | sb- | sb+ | sb- | sb+ | sb- |
| cross 1 | *gata2a<sup>+/+</sup>* | 16 | 1 | 20 | 0 | 0 | 0 | 23 | 4 |
|  | *gata2a<sup>um27/+</sup>* | 11 | 0 | 18 | 0 | 0 | 0 | 16 | 6 |
|  | *gata2a<sup>+/$\Delta$</sup>* | 0 | 0 | 0 | 0 | 1 | 0 | 7 | 1 |
|  | *gata2a<sup>um27/$\Delta$</sup>* | 0 | 0 | 0 | 0 | 0 | 1 | 0 | 21 |
| cross 2 | *gata2a<sup>+/+</sup>* | 24 | 0 | 25 | 0 | 0 | 0 | 22 | 3 |
|  | *gata2a<sup>um27/+</sup>* | 23 | 0 | 28 | 0 | 0 | 0 | 25 | 1 |
|  | *gata2a<sup>+/$\Delta$</sup>* | 0 | 0 | 0 | 0 | 0 | 0 | 34 | 3 |
|  | *gata2a<sup>um27/$\Delta$</sup>* | 0 | 0 | 0 | 0 | 0 | 1 | 0 | 20 |
| cross 3 | *gata2a<sup>+/+</sup>* | 22 | 1 | 18 | 1 | 0 | 0 | 23 | 1 |
|  | *gata2a<sup>um27/+</sup>* | 22 | 0 | 26 | 1 | 0 | 0 | 27 | 0 |
|  | *gata2a<sup>+/$\Delta$</sup>* | 0 | 0 | 0 | 0 | 1 | 0 | 22 | 3 |
|  | *gata2a<sup>um27/$\Delta$</sup>* | 0 | 0 | 0 | 0 | 0 | 0 | 1 | 22 |

1. *gata2a<sup>um27</sup>* was genotyped by KASP. Presence or deletion of exon 5 was detected by PCR. Exon 5 deletions ( $\Delta$ ) were only detected following injection with *cre* mRNA. *sb* – swim bladder, *cryG* – *cryaa:egfp* expression in lens

**Supplementary Table S2.** Phenotypic scoring of larvae from crosses between *gata2a^fl/fl^* and *gata2a^+/fl^* parents.

| <u>exon5 genotype<sup>1</sup></u> |  | uninjected |  |  |  | +25 pg zf<br>cre mRNA |  |  |  |
| --- | --- | --- | --- | --- | --- | --- | --- | --- | --- |
|  |  | cryG+ |  | cryG- |  | cryG+ |  | cryG- |  |
|  |  | sb+ | sb- | sb+ | sb- | sb+ | sb- | sb+ | sb- |
| cross 1 | *gata2a<sup>+/fl</sup>* | 21 | 0 | 0 | 0 | 0 | 0 | 0 | 0 |
|  | *gata2a<sup>fl/fl</sup>* | 15 | 0 | 0 | 0 | 0 | 0 | 0 | 0 |
|  | *gata2a<sup>+/$\Delta$</sup>* | 0 | 0 | 0 | 0 | 1 | 0 | 29 | 2 |
|  | *gata2a<sup>$\Delta/\Delta$</sup>* | 0 | 0 | 0 | 0 | 0 | 0 | 0 | 22 |
| cross 2 | *gata2a<sup>+/fl</sup>* | 16 | 0 | 0 | 0 | 0 | 0 | 0 | 0 |
|  | *gata2a<sup>fl/fl</sup>* | 20 | 0 | 0 | 0 | 0 | 0 | 0 | 0 |
|  | *gata2a<sup>+/$\Delta$</sup>* | 0 | 0 | 0 | 0 | 1 | 0 | 29 | 0 |
|  | *gata2a<sup>$\Delta/\Delta$</sup>* | 0 | 0 | 0 | 0 | 0 | 0 | 0 | 24 |
| cross 3 | *gata2a<sup>+/fl</sup>* | 11 | 0 | 0 | 0 | 0 | 0 | 0 | 0 |
|  | *gata2a<sup>fl/fl</sup>* | 11 | 0 | 0 | 0 | 0 | 0 | 0 | 0 |
|  | *gata2a<sup>+/$\Delta$</sup>* | 0 | 0 | 0 | 0 | 9 | 0 | 20 | 2 |
|  | *gata2a<sup>$\Delta/\Delta$</sup>* | 0 | 0 | 0 | 0 | 2 | 4 | 0 | 10 |

*sb* – swim bladder, *cryG* – *cryaa:egfp* expression in lens

**Supplementary Table 3.** Characterization of putative *foxc1a^fl^* carriers

|  |  |
| --- | --- |
| Total P0 screened | 45 |
| cryV<sup>+</sup>/mylC<sup>-</sup> only | 0 |
| cryV<sup>-</sup>/mylC<sup>+</sup> only | 0 |
| cryV<sup>+</sup>/mylC<sup>+</sup> only | 7 |
| 5'/3' PCR+ | 1** |
| 5'/3' PCR- | 6 |
| cryV<sup>+</sup>/mylC<sup>-</sup> &<br>cryV<sup>+</sup>/mylC<sup>+</sup> | 2 |
| 5'/3' PCR+ | 1* |
| 5'/3' PCR- | 1 |

\* - *foxc1a<sup>um383</sup>*
\*\* - cryV and mylC not genetically separable

## Notes

### Competing Interest Statement

The authors have declared no competing interest.

